# Cardiomyocyte BRAF is a key signalling intermediate in cardiac hypertrophy in mice

**DOI:** 10.1101/2022.09.07.506723

**Authors:** Hajed O. Alharbi, Michelle A. Hardyman, Joshua J. Cull, Thomais Markou, Susanna T.E. Cooper, Peter E. Glennon, Stephen J. Fuller, Peter H. Sugden, Angela Clerk

**Author notes:** **Correspondence to:** Angela Clerk,. These authors contributed equally (Hajed O. Alharbi, Michelle A. Hardyman).

## Abstract

Cardiac hypertrophy is necessary for the heart to accommodate an increase in workload. Physiological, compensated hypertrophy (e.g. with exercise) is reversible and largely due to cardiomyocyte hypertrophy. Pathological hypertrophy (e.g. with hypertension) is associated with additional features including increased fibrosis and can lead to heart failure. RAF kinases (ARAF/BRAF/RAF1) integrate signals into the ERK1/2 cascade, a pathway implicated in cardiac hypertrophy, and activation of BRAF in cardiomyocytes promotes compensated hypertrophy. Here, we used mice with tamoxifen-inducible cardiomyocyte-specific BRAF knockout (CM-BRAFKO) to assess the role of BRAF in hypertension-associated cardiac hypertrophy induced by angiotensin II (AngII; 0.8 mg/kg/d, 7 d) and physiological hypertrophy induced by phenylephrine (40 mg/kg/d, 7 d). Cardiac dimensions/function were assessed by echocardiography with histological assessment of cellular changes. AngII promoted cardiomyocyte hypertrophy and increased fibrosis within the myocardium (interstitial) and around the arterioles (perivascular) in male mice; cardiomyocyte hypertrophy and interstitial (but not perivascular) fibrosis were inhibited in mice with CM-BRAFKO. Phenylephrine had a limited effect on fibrosis, but promoted cardiomyocyte hypertrophy and increased contractility in male mice; cardiomyocyte hypertrophy was unaffected in mice with CM-BRAFKO, but the increase in contractility was suppressed and fibrosis increased. Phenylephrine induced a modest hypertrophic response in female mice and, in contrast to the males, tamoxifen-induced loss of cardiomyocyte BRAF reduced cardiomyocyte size, had no effect on fibrosis and increased contractility. The data identify BRAF as a key signalling intermediate in both physiological and pathological hypertrophy in male mice, and highlight the need for independent assessment of gene function in females.

**Clinical perspectives:**

- **Background**. BRAF is a key signalling intermediate that causes cancer and is upregulated in heart failure, but its role in physiological and pathological cardiac hypertrophy remains to be established.
- **Summary**. Cardiomyocyte BRAF is required in male mice for hypertrophy and contributes to interstitial fibrosis in hypertension induced by angiotensin II, but it increases contractility and suppresses fibrosis in physiological hypertrophy induced by α_1_-adrenergic receptor stimulation with phenylephrine. Differences between males and females are highlighted in the phenylephrine response.
- **Potential significance of results to human health and disease**. BRAF is a key signalling node in both pathological and physiological hypertrophy: inhibiting BRAF may be beneficial in pathological hypertrophy and the data have implications for repurposing of RAF inhibitors developed for cancer; inhibiting BRAF in physiological hypertrophy may result in increased fibrosis and using RAF inhibitors in this context could be detrimental in the longer term.

## Introduction

Cardiac hypertrophy is an important adaptive process required for the adult heart to accommodate an increase in workload. This may reflect “normal” life processes such as exercise or pregnancy, in which case the adaptation is largely reversible and described as physiological [1]. It is also necessary to accommodate stresses such as hypertension and this pathological hypertrophy is generally not reversible. Although the initial hypertrophy compensates for the developing disease, this can degenerate over time resulting in decompensation and heart failure. The adult mammalian heart is comprised largely of terminally-differentiated contractile cardiomyocytes with a network of capillaries that deliver blood throughout the heart from arterioles penetrating the myocardium. In addition, resident fibroblasts produce sufficient extracellular matrix for the heart to function. Irrespective of stimulus, the heart responds to an increased workload with hypertrophic growth of cardiomyocytes (in the absence of cell division) to increase contractile function [1]. In physiological hypertrophy (e.g. in response to exercise), this is beneficial and reversible, with little or no increase in fibrosis. In pathological conditions, cardiomyocyte hypertrophy is compromised by other pathological changes. These include loss of the capillary network and increased myocardial fibrosis, both of which potentially lead to cardiomyocyte dysfunction and cell death with a reduction in contractile ability of the heart. Understanding the molecular basis of these changes will aid in identifying therapeutic approaches for the treatment of heart failure, the main goals being to sustain cardiomyocyte function and survival, maintain the capillary network and prevent cardiac fibrosis [2].

A key signalling pathway linked to cardiac hypertrophy is the extracellular signal-regulated kinase 1/2 (ERK1/2) cascade. Activation of ERK1/2, the “original” mitogen-activated protein kinases (MAPKs), in cardiomyocytes promotes hypertrophic growth [3-5], but activation in cardiac fibroblasts promotes fibrosis [6, 7]. *In vivo* studies in mice show that mild constitutive-activation of the pathway in cardiomyocytes promotes compensated hypertrophy, whilst cardiomyocyte-specific gene deletion experiments indicate that ERK1/2 are not just important for hypertrophic growth, but are also cytoprotective [5, 8-11]. ERK1/2 are phosphorylated and activated by MAPK kinases 1/2 (MKK1/2) which are, in turn, phosphorylated by one or more of the RAF kinases (RAF1, BRAF, ARAF) [12]. RAF kinases form an integrating node into the cascade and are intimately linked to cancer. Indeed, oncogenic mutations in BRAF cause up to 30% of all cancers, and small molecule inhibitors of RAF kinases are already in use as anti-cancer drugs [13]. However, RAF kinases are not equivalent and their regulation is complex involving spatial organisation and interactions with multiple proteins, in addition to activating and inhibitory phosphorylations [12]. Importantly, although all RAF kinases can activate MKK1/2, ARAF and RAF1 have additional cytoprotective effects and inhibit pro-apoptotic kinases [14, 15].

Despite their importance, there are still relatively few studies of RAF kinases in cardiomyocytes and the heart. BRAF expression is upregulated in dilated cardiomyopathy and heart failure [16], suggesting it plays a role in human heart diseases. RAF1 and ARAF mRNAs are both downregulated in dilated cardiomyopathy, although there may be differential regulation of RAF1 in some forms of heart failure. All RAF kinases are expressed in rodent cardiomyocytes and are activated by hypertrophic stimuli such as endothelin-1 or α_1_-adrenergic receptor (α_1_-AR) agonists [17, 18]. *In vivo* studies in mice indicate that RAF1 is cytoprotective in cardiomyocytes, since overexpression of a dominant-negative form of RAF1 in cardiomyocytes or cardiomyocyte-specific RAF1 knockout leads to increased cardiomyocyte apoptosis and heart failure [19]. Activation of BRAF in cardiomyocytes by knock-in of the oncogenic V600E mutation promotes cardiomyocyte hypertrophy and compensated cardiac hypertrophy in mice [16] similar to that induced by overexpression of constitutively-active MKK1 [8]. Interestingly, RAF kinase inhibitors appear to have different effects on the heart. In a mouse model of hypertension induced by angiotensin II (AngII), dabrafenib inhibits both cardiomyocyte hypertrophy and cardiac fibrosis [20]. In contrast, encorafenib activates ERK1/2 signalling (via the RAF paradox [12]) and promotes compensated cardiac hypertrophy in a similar manner to knock-in of the BRAF(V600E) mutation [16]. These studies demonstrate that BRAF is important in cardiomyocytes and has the potential to drive cardiac and cardiomyocyte hypertrophy, but do not establish any involvement in cardiac hypertrophy, whether physiological or pathological.

Here, we developed a mouse model for inducible cardiomyocyte-specific knockout of BRAF. We show that BRAF is required for adaptation to hypertension, influencing both cardiomyocyte hypertrophy and fibrosis, but it plays a more subtle role affecting contractility and suppressing fibrosis in a model more akin to physiological hypertrophy. As with most studies, these experiments were conducted in male mice, but we also present data for physiological hypertrophy in female mice showing a different response to the males. These data highlight the need for full and proper experimental assessment of the underlying mechanisms of cardiac hypertrophy in females.

## Methods

### Ethics statement for animal experiments

Mice were housed at the BioResource Unit at University of Reading or St. George’s University of London (UK registered with a Home Office certificate of designation). All procedures were performed in accordance with UK regulations and the European Parliament Directive 2010/63/EU for animal experiments. Work was undertaken in accordance with local institutional animal care committee procedures at the University of Reading and the U.K. Animals (Scientific Procedures) Act 1986. Studies were conducted under Project Licences 70/8248, 70/8249 and P8BAB0744.

### *In vivo* studies of mice with cardiomyocyte-specific deletion of BRAF

Housing conditions were as described in [16, 21]. Breeding was conducted with mice between 6 weeks and 8 months with a maximum of 4 litters per female. Mice undergoing procedures were monitored using a score sheet and routinely culled if they reached a predefined endpoint agreed with the Named Veterinary Surgeon. Weights were taken before, during and at the end of the procedures. Mouse weights from the start and end of procedures are provided in **Supplementary Table S1**. Mice were allocated to specific groups on a random basis with randomisation performed independently of the individual leading the experiment. No mice were excluded after randomisation. Individuals conducting the studies were not blinded to experimental conditions for welfare monitoring purposes. Data and sample analysis (e.g. echocardiography, histology) was performed by individuals who were blinded to intervention.

#### Mouse lines

Genetically-modified mice were from Jackson Laboratories, imported into the UK and transported to University of Reading for breeding in-house. We used mice with a floxed cassette for Cre-induced BRAF gene deletion (129-*Braf*^*tm1Sva*^/J, strain no. 006373) [22]. Mice were backcrossed against the C57Bl/6J background at University of Reading for at least 4 generations prior to experimentation. Mice were bred with *Myh6*-MERCreMER mice expressing tamoxifen-inducible Cre recombinase under control of a mouse *Myh6* promoter [Tg(Myh6-cre)1Jmk/J, strain no. 009074] [23]. Breeding protocols were used to produce male and female BRAF^fl/fl^/Cre^+/-^ mice (i.e. homozygous for floxed BRAF and hemizygous for Cre) for experimentation. Additional studies were conducted in parallel with wild-type mice from the same breeding stocks.

#### Genotyping and confirmation of recombination

DNA was extracted from ear clips (taken for identification purposes) using Purelink genomic DNA (gDNA) mini-kits (Invitrogen). Briefly, tissue was digested in genomic digestion buffer containing proteinase K (overnight, 55°C). Following centrifugation (12,000 × g, 3 min, 18°C), supernatants were incubated with RNAse A (2 min) before addition of genomic lysis binding buffer mixed with an equal volume of ethanol. gDNA was purified using Purelink spin columns and PCR amplified with specific primers (see **Supplementary Table S2** for primer sequences and annealing temperatures) using GoTaq Hot Start Polymerase (Promega). PCR conditions were 95°C for 3 min, followed by up to 33 cycles of 95°C denaturation for 30 s, 30 s annealing, elongation at 72°C for 30 s, followed by a 7 min 72°C final extension. PCR products were separated on 2% (w/v) agarose gels (25 min, 80 V) and visualised under UV light. Mice (males: 7-8 weeks; females: 9-10 weeks) were treated with a single dose of tamoxifen (40 mg/kg i.p.; Sigma-Aldrich) or vehicle alone. Tamoxifen was dissolved in 0.25 ml ethanol and then mixed with 4.75 ml corn oil. For confirmation of recombination, RNA was extracted from tissue powders and cDNA prepared as described below. cDNA (4 µl) was subjected to PCR analysis using GoTaq Hot Start Polymerase with specific primers and conditions (see **Supplementary Table S2** for primer sequences and annealing temperatures). PCR conditions were 95°C for 3 min, followed by 32 cycles of 95°C denaturation for 30 s, 30 s at 57°C, elongation at 72°C for 60 s, followed by a 7 min 72°C final extension. Products were separated by electrophoresis on a 2% (w/v) agarose gel (85 V, 45 min).

#### Drug delivery to induce cardiac hypertrophy

Drug delivery used Alzet osmotic pumps (model 1007D; supplied by Charles River), filled according to the manufacturer’s instructions in a laminar flow hood using sterile technique. Mice were treated with 0.8 mg/kg/d angiotensin II (AngII) in acidified PBS (PBS containing 10 mM acetic acid) or with acidified PBS alone as in [20, 24]. We used 0.8 mg/kg/d AngII as a moderate concentration that induces hypertension [25] rather than a subpressor dose (e.g. 0.288 mg/kg/d [26]) or a high dose that can be associated with sudden cardiac death (e.g. >2 mg/kg/d [7]).

Alternatively, mice were treated with 40 mg/kg/d phenylephrine in PBS or with PBS alone as in [27]. Minipumps were incubated overnight in sterile PBS (37°C). Mice were given 0.05 mg/kg (s.c.) buprenorphine (Vetergesic, Ceva Animal Health Ltd.) before being placed under continuous inhalation anaesthesia using isoflurane (induction at 5%, maintenance at 2 - 2.5%) mixed with 2 l/min O_2_. A 1 cm incision was made in the mid-scapular region and minipumps were implanted portal first in a pocket created in the left flank region of the mouse. Wound closure used wound clips or a simple interrupted suture with polypropylene 4-0 thread (Prolene, Ethicon). Mice were recovered singly and returned to their home cage once fully recovered.

#### Cardiac ultrasound

Echocardiography was performed using a Vevo 2100 equipped with a MS400 18-38 MHz transducer (Visualsonics) as in [16, 20, 27]. Mice were anaesthetised in an induction chamber with isoflurane (5% flow rate) with 1 l/min O_2_ and transferred to the heated Vevo Imaging Station. Anaesthesia was maintained with 1.5% isoflurane delivered via a nose cone. Left ventricular cardiac function and structure was assessed from short axis M-mode images with the axis placed at the mid-level of the left ventricle at the level of the papillary muscles. Left ventricular mass and chamber volumes were measured from B-mode long axis images using VevoStrain software for speckle tracking. Baseline scans were taken prior to experimentation (−7 to −3 days). Further scans were taken at intervals following minipump implantation. Imaging was completed within 20 min. Data analysis (Vevo LAB) was performed by an independent assessor blinded to intervention. Data were gathered from two scans taken from each time point, taking mean values across 4 cardiac cycles for each M-mode scan or 2 cardiac cycles for long axis B-mode images. Mice were recovered singly and transferred to the home cage once fully recovered.

#### Tissue harvesting and histology

Mice were culled by CO_2_ inhalation followed by cervical dislocation. Hearts were excised quickly, washed in PBS, dried and snap-frozen in liquid N_2_ or fixed for histology. Histological sections were prepared and stained by HistologiX Limited. Haemotoxylin and eosin staining was used for analysis of myocyte cross-sectional area. Cells around the periphery of the left ventricle (excluding epicardial layer) were chosen at random and outline traced using NDP.view2 software (Hamamatsu). Up to 30 cells were measured per section by a single independent assessor and the mean value taken for each mouse. For analysis of fibrosis, we used picrosirius red staining and the whole section was scored for perivascular fibrosis around arterioles (identified by a clear elastic layer) and interstitial fibrosis. For perivascular fibrosis values were from 1 (negligible increase in fibrosis around any vessel), through to 4 (extensive fibrosis around multiple vessels, penetrating into the myocardium). For interstitial fibrosis values were from 1 (negligible increase in fibrosis within the myocardium distal from any arterioles), through to 4 (extensive and pervasive fibrosis throughout the left ventricle).

### RNA preparation and qPCR

Mouse heart powders were weighed into safelock eppendorf tubes and kept on dry ice. RNA Bee (AMS Biotechnology Ltd) was added (1 ml per 10-15 mg) and the samples homogenised on ice using a pestle. RNA was prepared according to the manufacturer’s instructions and dissolved in nuclease-free water. The purity was assessed from the A_260_/A_280_ measured using an Implen NanoPhotometer (values were 1.8–2.0) and concentrations determined from the A_260_. Quantitative PCR (qPCR) analysis was performed as described in [28]. Total RNA (1 µg) was reverse transcribed to cDNA using High Capacity cDNA Reverse Transcription Kits with random primers (Applied Biosystems). qPCR was performed using an ABI Real-Time PCR 7500 system (Applied Biosystems) using 1/40 of the cDNA produced. Optical 96-well reaction plates were used with iTaq Universal SYBR Green Supermix (Bio-Rad Laboratories Inc.) according to the manufacturer’s instructions. See **Supplementary Table S3** for primer sequences. Results were normalized to *Gapdh*, and relative quantification was obtained using the ΔCt (threshold cycle) method; relative expression was calculated as 2^−ΔΔCt^, and normalised as indicated in the Figure Legends.

### Immunoblotting

Heart powders (15-20 mg) were extracted in 6 vol extraction buffer [20 mM Tris pH 7.5, 1 mM EDTA, 10% (v/v) glycerol, 1% (v/v) Triton X-100, 100 mM KCl, 5 mM NaF, 0.2 mM Na_3_VO_4_, 5 mM MgCl_2_, 0.05% (v/v) 2-mercaptoethanol, 10 mM benzamidine, 0.2 mM leupeptin, 0.01 mM trans-epoxy succinyl-l-leucylamido-(4-guanidino)butane, 0.3 mM phenylmethylsulphonyl fluoride, 4 µM microcystin]. Samples were vortexed and extracted on ice (10 min), then centrifuged (10,000 × g, 10 min, 4°C). The supernatants were removed, a sample was taken for protein assay and the rest boiled with 0.33 vol sample buffer (300 mM Tris-HCl pH 6.8, 10% (w/v) SDS, 13% (v/v) glycerol, 130 mM dithiothreitol, 0.2% (w/v) bromophenol blue). Protein concentrations were determined by BioRad Bradford assay using BSA standards.

Proteins were separated by SDS-PAGE (200 V) using 8% (for RAF kinases), or 12% (Gapdh) polyacrylamide resolving gels with 6% stacking gels until the dye front reached the bottom of the gel (∼50 min). Proteins were transferred electrophoretically to nitrocellulose using a BioRad semi-dry transfer cell (10 V, 60 min). Non-specific binding sites were blocked (15 min) with 5% (w/v) non-fat milk powder in Tris-buffered saline (20 mM Tris-HCl pH 7.5, 137 mM NaCl) containing 0.1% (v/v) Tween 20 (TBST). Blots were incubated with primary antibodies in TBST containing 5% (w/v) BSA (overnight, 4°C), then washed with TBST (3 × 5 min, 21°C), incubated with horseradish peroxidase-conjugated secondary antibodies in TBST containing 1% (w/v) non-fat milk powder (60 min, 21°C) and then washed again in TBST (3 × 5 min, 21°C). Antibodies to RAF1 were from BD Transduction Labs (mouse monoclonal, Cat. No. 610152), antibodies to BRAF and ARAF were from Santa Cruz Biotechnology Inc. (BRAF: mouse monoclonal, Cat. No. sc-5284; ARAF: rabbit polyclonal, Cat. No. sc-408), antibodies to Gapdh were from Cell Signaling Technologies (rabbit monoclonal, Cat. No. 5174) and secondary antibodies were from Dako, supplied by Agilent (rabbit anti-mouse immunoglobulins/HRP, Cat. No. P0260; Goat anti-rabbit immunoglobulins/HRP, Cat. No. P0448). Primary antibodies were used at 1/1000 dilustion. Secondary antibodies were used at 1/5000 dilution.

Bands were detected by enhanced chemiluminescence using ECL Prime with visualisation using an ImageQuant LAS4000 system (Cytiva). ImageQuant TL 8.1 software (GE Healthcare) was used for densitometric analysis. Raw values for phosphorylated kinases were normalised to the total kinase. Values for all samples were normalised to the mean of the controls.

### Image processing and statistical analysis

Images were exported from the original software as .tif or .jpg files and cropped for presentation using Adobe Photoshop CC maintaining the original relative proportions. Data analysis used Microsoft Excel and GraphPad Prism 9. Statistical analysis was performed using GraphPad Prism 9 with two-tailed unpaired t tests, or two-tailed one-way or two-way ANOVA as indicated in the Figure Legends. A multiple comparison test was used in combination with ANOVA. A Grubb’s outlier test was applied to the data, and outliers excluded from the analysis. Graphs were plotted with GraphPad Prism 9. Specific p values are provided with significance levels of p<0.05 in bold type.

## Results

### Cardiomyocyte BRAF knockout model

Our previous studies demonstrated that activation of BRAF in cardiomyocytes can promote hypertrophy [16], but this does not establish the role it may play in adaptation of the heart to pathophysiological stresses. To assess this, we generated mice for cardiomyocyte-specific inducible BRAF knockout by crossing mice with a floxed cassette for Cre-induced BRAF gene deletion [22] with *Myh6*-MERCreMER mice expressing tamoxifen-inducible Cre recombinase under control of a mouse *Myh6* promoter [23]. Experiments first used male mice homozygous for floxed BRAF and hemizygous for Cre (BRAF^fl/fl^/Cre^+/-^ mice). Baseline echocardiograms were collected at age 7-8 weeks, then cardiomyocyte BRAF knockout was induced by a single injection of tamoxifen (40 mg/kg i.p.). Recombination was confirmed at the mRNA level and decreased expression of BRAF in the heart was confirmed by immunoblotting (**Figure 1A,B**). RAF1 (not ARAF) expression was also decreased. This may be because BRAF forms heterodimers with RAF1 in cardiomyocytes [16], so loss of BRAF potentially affects expression of the pool of RAF1 with which it associates. Tamoxifen and cardiomyocyte BRAF knockout alone had no significant effects on any of the parameters studied relative to BRAF^fl/fl^/Cre^+/-^ mice treated with corn-oil vehicle (e.g. hypertrophic gene markers, **Figure 1C**), indicating that BRAF is not essential in normal adult mouse hearts, at least in the short term. BRAF^fl/fl^/Cre^+/-^ mice exhibited no baseline differences in heart morphology compared with wild-type C57Bl/6J mice from the same breeding stocks studied in parallel, with similar sized cardiomyocytes and little fibrosis (**Figure 1D**). The hypertrophic response in BRAF^fl/fl^/Cre^+/-^ mice induced by AngII was also similar to wild-type mice (**Figure 1E**). In summary, we detected no significant baseline effects of the mutant genetic background in male mice with or without cardiomyocyte BRAF knockout.

**Figure 1.**
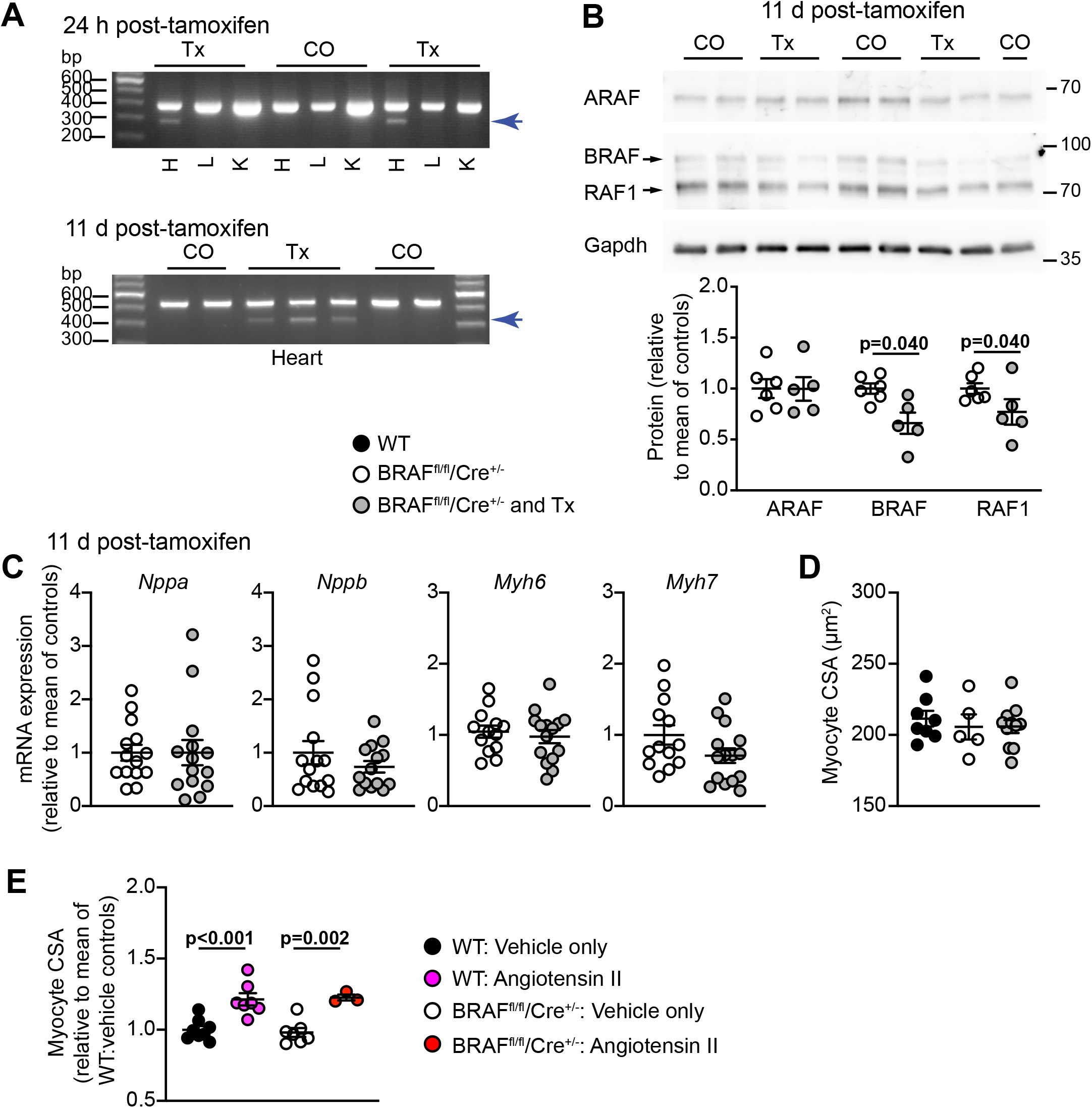
Male mice with tamoxifen-inducible cardiomyocyte-specific BRAF knockout. BRAF^fl/fl^/Cre^+/-^ (homozygous for floxed BRAF; hemizygous for Cre) male mice were treated with corn-oil (CO) or tamoxifen in corn-oil (Tx) for 24 h or 11 d as indicated. **A**, Confirmation of BRAF knockout using cDNA prepared from RNA extracted from powdered tissues. PCR amplification used forward primers in exon 10 (upper image) or exon 9 (lower image) with reverse primers in exon 13. Deletion of exon 12 in cardiomyocytes resulted in the appearance of a smaller product following recombination with tamoxifen administration in heart (H), but not liver (L) or kidney (K). Representative images are shown. **B**, Immunoblot analysis of RAF isoforms in samples of mouse hearts treated with vehicle or tamoxifen. Representative immunoblots are in the upper panels (positions of relative molecular mass markers are on the right) with densitometric analysis below. RAF expression was normalised to Gapdh and data presented relative to the means of the vehicle treated controls. Statistical analysis used unpaired two-tailed t tests. **C**, mRNA expression in mouse hearts after 11 d treatment with tamoxifen. RNA was extracted and expression of selected genes assessed by qPCR. **D**,**E**, Assessment of cardiomyocyte cross-sectional area (CSA) in wild-type (WT) mice from the same breeding stock as the BRAF^fl/fl^/Cre^+/-^ mice in comparison with the BRAF^fl/fl^/Cre^+/-^ mice. Mice were treated with Tx for 11 d, angiotensin II (0.8 mg/kg/d, 7 d) in acidified PBS (AcPBS) or AcPBS only. Data are from haematoxylin & eosin stained sections. Data are presented as individual values with means ± SEM. Statistical analysis used unpaired two-tailed t tests.

### Cardiomyocyte BRAF is required for cardiac adaptation to hypertension induced by angiotensin II

We assessed the effects of BRAF knockout in a model of hypertension induced by treatment with 0.8 mg/kg/d AngII over 7 d (**Figure 2A**). AngII causes pressure-overload on the heart by increasing systemic blood pressure and stimulates endothelial cells lining the blood vessels. The primary impact on the heart is mechanical stress, first on the arteries/arterioles and, subsequently, on the rest of the myocardium. AngII in acidified PBS (AcPBS) or AcPBS alone was delivered by osmotic minipumps, implanted 4 d after tamoxifen administration, by which time tamoxifen has cleared the body [29]. As in previous studies [20, 24], AngII promoted cardiac hypertrophy: left ventricular (LV) internal diameter and predicted end diastolic volume decreased, whilst LV wall thickness and predicted end diastolic LV mass were increased (**Figure 2B.C; Supplementary Table S4**). At 7 d, we detected no significant change in ejection fraction or fractional shortening (**Supplementary Table S4**). Cardiomyocyte-specific deletion of BRAF did not affect the decrease in LV internal diameter or volume, but reduced AngII-induced increases in LV wall thickness and predicted mass (**Figure 2B,C**). AngII increased cardiomyocyte cross-sectional area (a measure of hypertrophy) to a similar degree in wild-type and BRAF^fl/fl^/Cre^+/-^ mice, and this was inhibited by cardiomyocyte BRAF knockout (**Figures 1E and 2D**).

**Figure 2.**
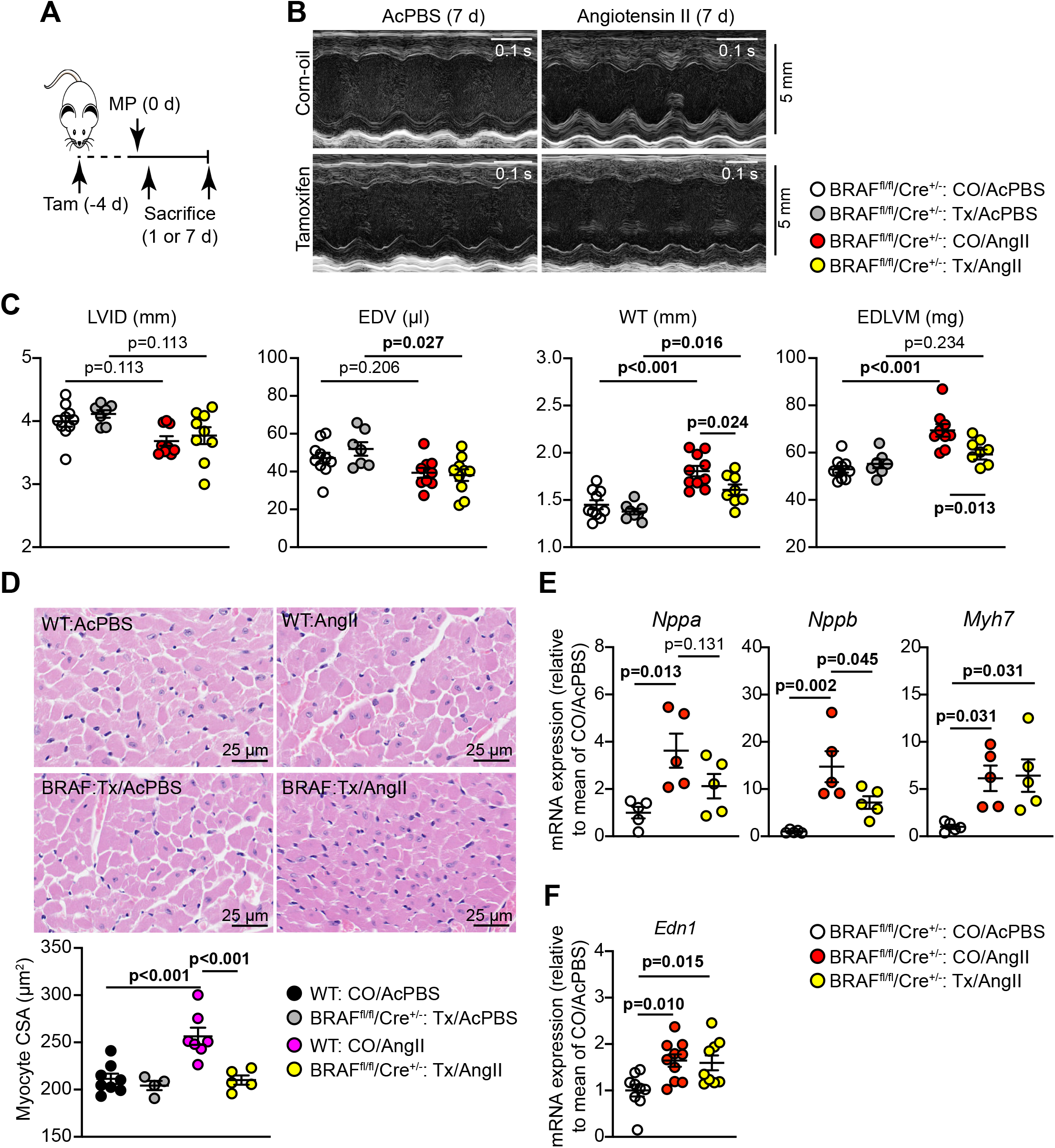
Cardiomyocyte BRAF knockout inhibits the hypertrophic response induced by angiotensin II in male mouse hearts. Male BRAF^fl/fl^/Cre^+/-^ (BRAF) or wild-type (WT) mice were treated with corn-oil vehicle (CO) or tamoxifen in corn-oil (Tx) 4 days before minipumps (MP) were implanted to deliver acidified PBS (AcPBS) or 0.8 mg/kg/d angiotensin II (AngII). Mice were sacrificed after 1 or 7 d. Echocardiograms were collected and hearts taken at 7 d. **A**, Schematic of experimental protocol. **B**, Representative M-mode echocardiograms taken from short axis views of the heart at 7 d. **C**, Analysis of echocardiograms taken at 7 d to assess cardiac dimensions. LVID, left ventricle internal diameter; EDV, end diastolic volume; WT, left ventricle wall thickness (posterior plus anterior walls); EDLVM, end diastolic left ventricular mass. LVID and WT were measured at diastole from M-mode images of short axis views of the heart. EDV and EDLVM were predicted from B-mode images of long axis views of the heart. **D**, Haematoxylin and eosin staining of mouse heart sections (upper panels) from hearts collected at 7 d, with assessment of cardiomyocyte cross-sectional area (CSA; lower panel). Images and measurements are from the periphery of the left ventricle. **E-F**, mRNA expression in mouse hearts after 24 h (**E**) or 7 d (**F**) treatment with angiotensin II. RNA was extracted and expression of *Nppa, Nppb, Myh7* and *Edn1* were assessed by qPCR. Data are individual values with means ± SEM. Statistical analysis used two-way (**C**,**D**) or one-way (**E**,**F**) ANOVA with Holm-Sidak’s post-test. Statistically significant values (p<0.05) are in bold type.

Pathological hypertrophy is associated with increases in cardiac mRNA expression of the “foetal” gene markers, *Nppa, Nppb*, and *Myh7*. These markers were not significantly changed with cardiomyocyte BRAF knockout alone (**Figure 1C**), but AngII-induced increases in expression of *Nppa* and *Nppb* (not *Myh7*) were inhibited in mouse hearts with cardiomyocyte deletion of BRAF (**Figure 2E**). As mentioned, AngII causes pressure-overload on the heart by increasing systemic blood pressure. Cardiomyocyte hypertrophy may be stimulated by biomechanical stresses detected by mechanosensors in the cell (e.g. stretch-regulated ion channels [30] or the myofibrillar apparatus and other structural components [31]) that trigger changes in gene expression and cause hypertrophy. Additionally, neurohumoral mediators produced by the vessels may contribute. For example, endothelin-1 produced by endothelial cells in response to AngII [32] is a potent stimulus of cardiomyocyte hypertrophy, signalling through the ERK1/2 pathway [3]. Notably, endothelin-1 mRNA (*Edn1*) was upregulated by AngII, a response that was unaffected by cardiomyocyte BRAF knockout (**Figure 2F**). Irrespective of the origin of the signal, cardiomyocyte BRAF is a key mediator of cardiomyocyte hypertrophy induced by AngII.

Cardiac hypertrophy is associated with cardiomyocyte growth, but can also reflect increased fibrosis. This may be mediated via ERK1/2 signalling, and influenced by cardiomyocytes acting on neighbouring cells (e.g. via paracrine mediators) [33-35]. AngII increased cardiac fibrosis (shown with picrosirius red staining), with a striking effect in the immediate environment of the arterioles within the myocardium (the “perivascular” region; **Figure 3A-C**). AngII promoted a relatively small increase in interstitial fibrosis and this was generally in localised areas of the heart. Interstitial fibrosis, but not perivascular fibrosis, was reduced in hearts with cardiomyocyte BRAF knockout (**Figure 3C**). Thus, interstitial and perivascular fibrosis have independent origins and, whilst cardiomyocyte hypertrophy contributes to the accumulation of interstitial fibrosis, the impact of hypertension on the fibrosis surrounding arterioles is largely independent of cardiomyocyte involvement. The data are consistent with our observation that upregulation of *Edn1* mRNA was not inhibited by cardiomyocyte BRAF knockout (**Figure 2F**) since mediators deriving from the vessels would not be expected to be affected. In contrast, accumulation of fibrotic material within the myocardium is potentially driven by the cardiomyocytes themselves. In support of this, *Fgf2* mRNA expression was significantly upregulated in hearts of mice treated with AngII, and the increase in expression was reduced in mouse hearts with cardiomyocyte BRAF knockout (**Figure 3D)**. This is consistent with the concept of cross-talk between cardiomyocytes and the non-myocytes in the heart [36].

**Figure 3.**
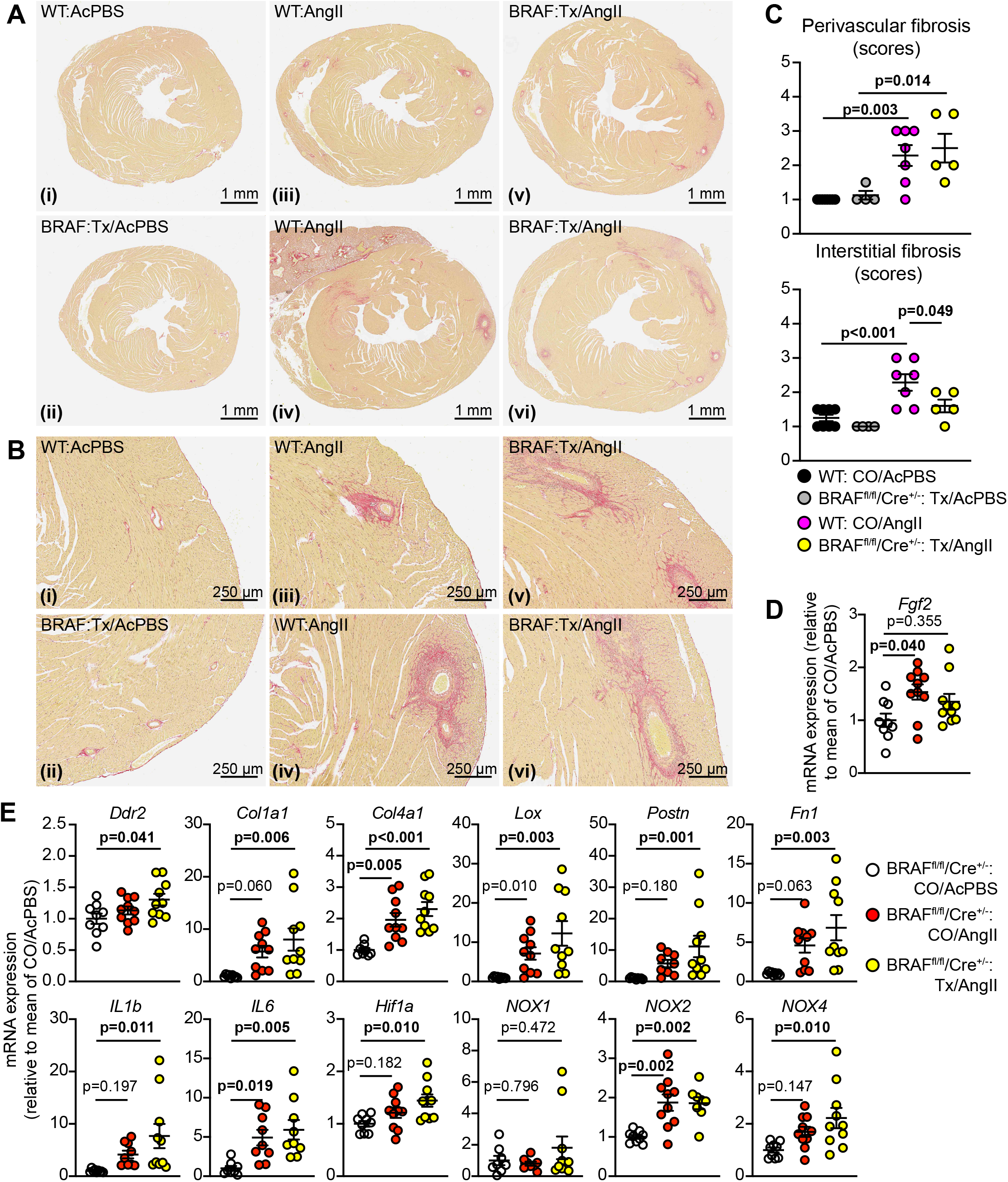
Cardiomyocyte BRAF knockout inhibits interstitial but not perivascular fibrosis induced by angiotensin II in male mouse hearts. Male BRAF^fl/fl^/Cre^+/-^ (BRAF) or wild-type (WT) mice were treated with corn-oil (CO) or tamoxifen in corn-oil (Tx) 4 days before minipumps were implanted to deliver acidified PBS (AcPBS) or 0.8 mg/kg/d angiotensin II in AcPBS (AngII) for 7 d. **A**,**B**, Picrosirius red staining of mouse heart sections showing short axis views of the whole heart (**A**) with enlarged sections (**B**) from the same views (the red stain shows accumulation of fibrotic material). Representative (average) images are shown for mice treated with CO/AcPBS (i), Tx/AcPBS (ii), CO/AngII (iii) and Tx/AngII (v). Additional images are shown for the most severe degree of fibrosis with CO/AngII (iv) and Tx/AngII (vi). **C**, Quantification of the degree of fibrosis. This was scored as: 1 = the least amount of fibrosis; 2 = low level fibrosis; 3 = high level fibrosis in at least one area of the myocardium; 4 = high level fibrosis throughout the myocardium (half scores were used). **D**,**E**, mRNA expression in mouse hearts. RNA was extracted and expression of selected genes assessed by qPCR. Data are presented as individual values with means ± SEM. Statistical analysis used two-way (**C**) or one-way (**D**,**E**) ANOVA with Holm-Sidak’s post-test. Statistically significant values (p<0.05) are in bold type.

To gain further insight into the consequences of cardiomyocyte BRAF knockout for the heart in AngII-induced cardiac hypertrophy, we assessed mRNA expression of selected genes. Despite the effect on *Fgf2* (**Figure 3D**), upregulation of fibrotic gene markers by AngII was generally enhanced in hearts from mice with cardiomyocyte BRAF knockout (**Figure 3E**), including the fibroblast marker *Ddr2*, collagens (*Col1a1, Col1a2*), other extracellular matrix proteins (*Postn, Fn1*) and lysyl oxidase (*Lox*) involved in collagen cross linking. This presumably reflects the prevalent fibrosis in the perivascular region (**Figure 3A-C**). Other genes associated with hypoxia (*Hif1a*) and inflammation (*IL1b, IL6*) were also enhanced in cardiomyocyte BRAF knockout hearts (**Figure 3E**) and may contribute to further development of fibrosis [33]. AngII-induced hypertrophy is associated with increased reactive oxygen species (ROS), potentially from NADPH oxidases NOX2 and NOX4 [37]. Consistent with other studies, NOX2 mRNA expression was increased with AngII and there was some increase in NOX4 (though not NOX1). Only NOX4 expression was further enhanced by cardiomyocyte BRAF knockout and, as a constitutively active enzyme, this is predicted to increase cardiac oxidative stress [33, 37]. Overall, these changes in gene expression indicate that the pathophysiological stresses resulting from AngII-induced hypertension were not ameliorated in the hearts with cardiomyocyte BRAF knockout and reduced cardiomyocyte hypertrophy and were, if anything, exacerbated.

### Cardiomyocyte BRAF does not drive cardiomyocyte hypertrophy in response to phenylephrine and suppresses fibrosis in male mice

In contrast to AngII, phenylephrine (as an α_1_-AR agonist) is associated with adaptive physiological hypertrophy and, although there are additional systemic effects, it promotes compensated cardiac hypertrophy acting directly on the cardiomyocytes [38]. Male mice were treated with 40 mg/kg/d phenylephrine in PBS or PBS alone (7 d) using osmotic minipumps, implanted 4 d after tamoxifen administration as for AngII (**Figure 2A**). As in previous studies [27], phenylephrine promoted cardiac hypertrophy with decreased LV internal diameter and increased wall thickness, and neither appeared to be significantly affected in hearts of mice with cardiomyocyte BRAF knockout (**Figure 4A; Supplementary Table S5**). Furthermore, the increase in cardiomyocyte cross-sectional area induced by phenylephrine was not significantly reduced by cardiomyocyte BRAF knockout (**Figure 4B**). This is consistent with our previous studies showing that phenylephrine promotes cardiomyocyte hypertrophy via insulin receptor family members and protein kinase B (PKB, or Akt) [27]. Phenylephrine alone promoted relatively little increase in cardiac fibrosis (**Figure 4C,D**), particularly compared with AngII (**Figure 3A,B**). Surprisingly, both perivascular and interstitial fibrosis were significantly and substantially increased in hearts of mice with cardiomyocyte BRAF knockout when treated with phenylephrine, with a particularly striking effect on interstitial fibrosis that in some cases permeated the full cross-sectional area of the myocardium (**Figure 4C-E**). As with AngII (**Figure 3A-C**), the effects of cardiomyocyte BRAF knockout on “physiological” hypertrophy induced by phenylephrine (**Figure 4C-E**) suggest that the origin of interstitial fibrosis is independent of perivascular fibrosis and controlled to some degree by the cardiomyocytes.

**Figure 4.**
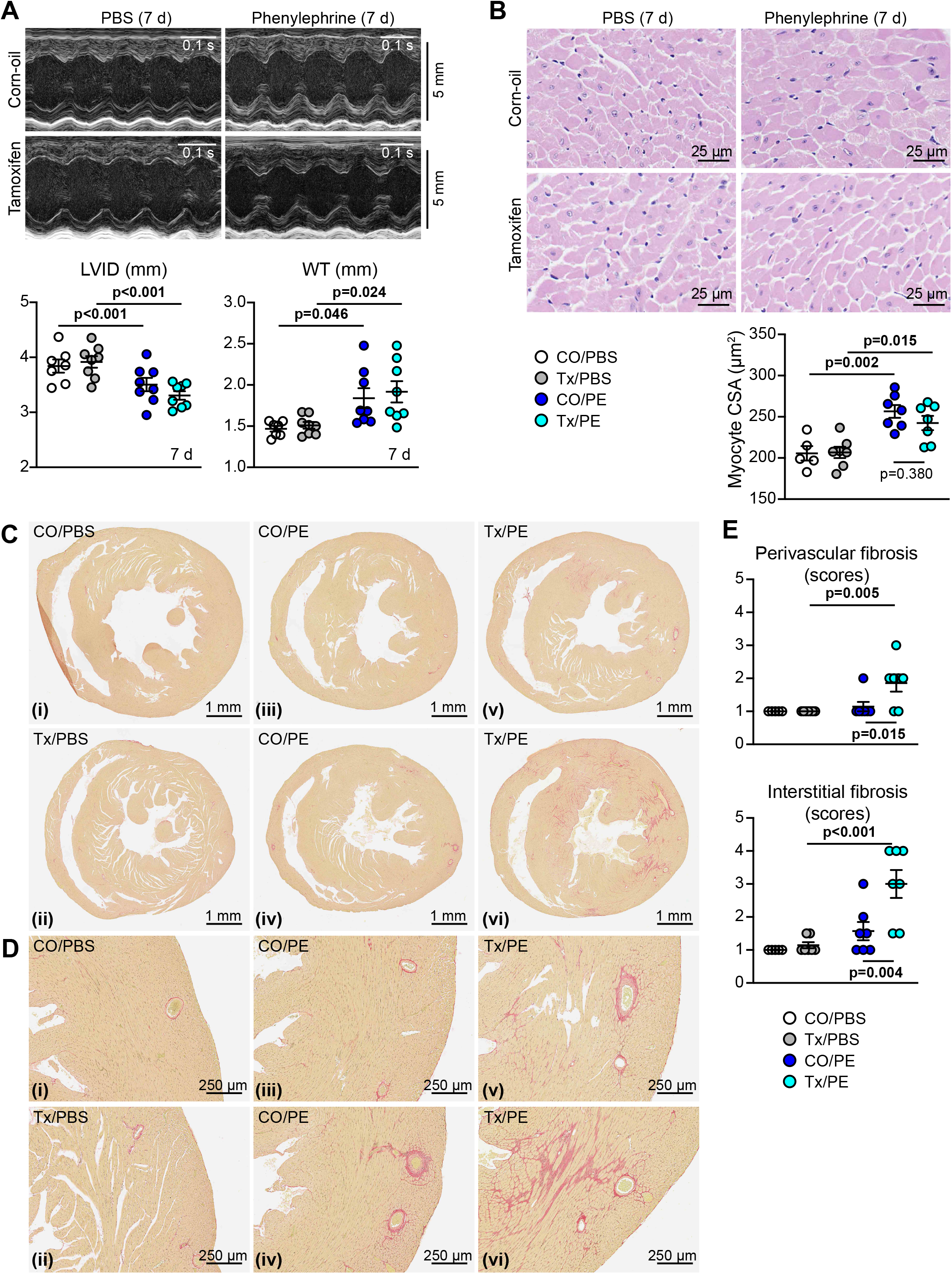
Cardiomyocyte BRAF knockout does not inhibit cardiomyocyte hypertrophy induced by phenylephrine in male mouse hearts, but increases cardiac fibrosis. Male BRAF^fl/fl^/Cre^+/-^ mice were treated with corn-oil (CO) or tamoxifen in corn-oil (Tx) 4 days before minipumps were implanted to deliver PBS or 40 mg/kg/d phenylephrine in PBS (PE) for 7 d. **A**, Representative M-mode echocardiograms taken from short axis views of the heart (upper panels) with analysis of echocardiograms to assess cardiac dimensions (lower panels). LVID, left ventricle internal diameter; WT, left ventricle wall thickness (posterior plus anterior walls). Diastolic measurements are shown. **B**, Haematoxylin and eosin staining of mouse heart sections (upper panels) with assessment of cardiomyocyte cross-sectional area (CSA; lower panel). Images and measurements are from the periphery of the left ventricle. **C**,**D**, Picrosirius red staining of mouse heart sections showing short axis views of the whole heart (**C**) with enlarged sections (**D**) from the same views (the red stain shows accumulation of fibrotic material). Representative (average) images are shown for mice treated with CO/PBS (i), Tx/PBS (ii), CO/PE (iii) and Tx/PE (v). Additional images are shown for the most severe degree of fibrosis with CO/PE (iv) and Tx/PE (vi). **E**, Quantification of fibrosis. This was scored as: 1 = the least amount of fibrosis; 2 = low level fibrosis; 3 = high level fibrosis in at least one area of the myocardium; 4 = high level fibrosis throughout the myocardium (half scores were used). Data are presented as individual values with means ± SEM. Statistical analysis used two-way ANOVA with Holm-Sidak’s post-test. Statistically significant values (p<0.05) are in bold type.

### Cardiomyocyte BRAF knockout in female versus male mice: effects on phenylephrine-induced cardiac hypertrophy

There are sex-specific differences in the physiological hypertrophic response of the heart in female vs male animals (e.g. in exercise-induced hypertrophy [39]), probably due in part to sex hormones and influences on metabolism [40]. We therefore assessed the role of cardiomyocyte BRAF in the hypertrophic response to phenylephrine in the female BRAF^fl/fl^/Cre^+/-^ littermates of the males used in the experiments outlined above. Because of their smaller size, experiments were initiated with female mice at 9-10 weeks of age. The females were still significantly smaller than the males, but (as with the males) there was no significant difference in body weights between the groups of mice with different treatments at baseline or at the end of the experiment (**Figure 5A; Supplementary Table S1**). Mice were anaesthetized for echocardiography and cardiac function and dimensions were assessed prior to intervention. We detected no significant difference in heart rate, ejection fraction or fractional shortening between male and female hearts, nor was there any difference in global longitudinal or circumferential strain (**Figure 5B**). Thus, cardiac function was similar in males and females. There was a small, but significant decrease in stroke volume and cardiac output, probably because of the small (non-significant) decrease in diastolic LV internal diameter and predicted end diastolic volume, in the absence of any difference in systolic measurements (**Figure 5C**). However, predicted end diastolic LV mass was ∼13.8% greater in male mice than in female mice [49.56 ± 0.87 mg (n=31) and 43.56 ± 0.77 mg (n=39), respectively (means ± SEM)] (**Figure 5D**). This was reflected in the significant difference in diastolic and systolic LV wall thickness. There was no evidence for recombination in female or male hearts in the absence of tamoxifen, and treatment with tamoxifen induced a similar degree of recombination (and therefore cardiomyocyte BRAF knockout) in females as in males (**Figure 5E**). Treatment with tamoxifen resulted in a decrease in BRAF protein but, unlike the males, we did not detect any decrease in RAF1 protein (**Figure 5F**).

**Figure 5.**
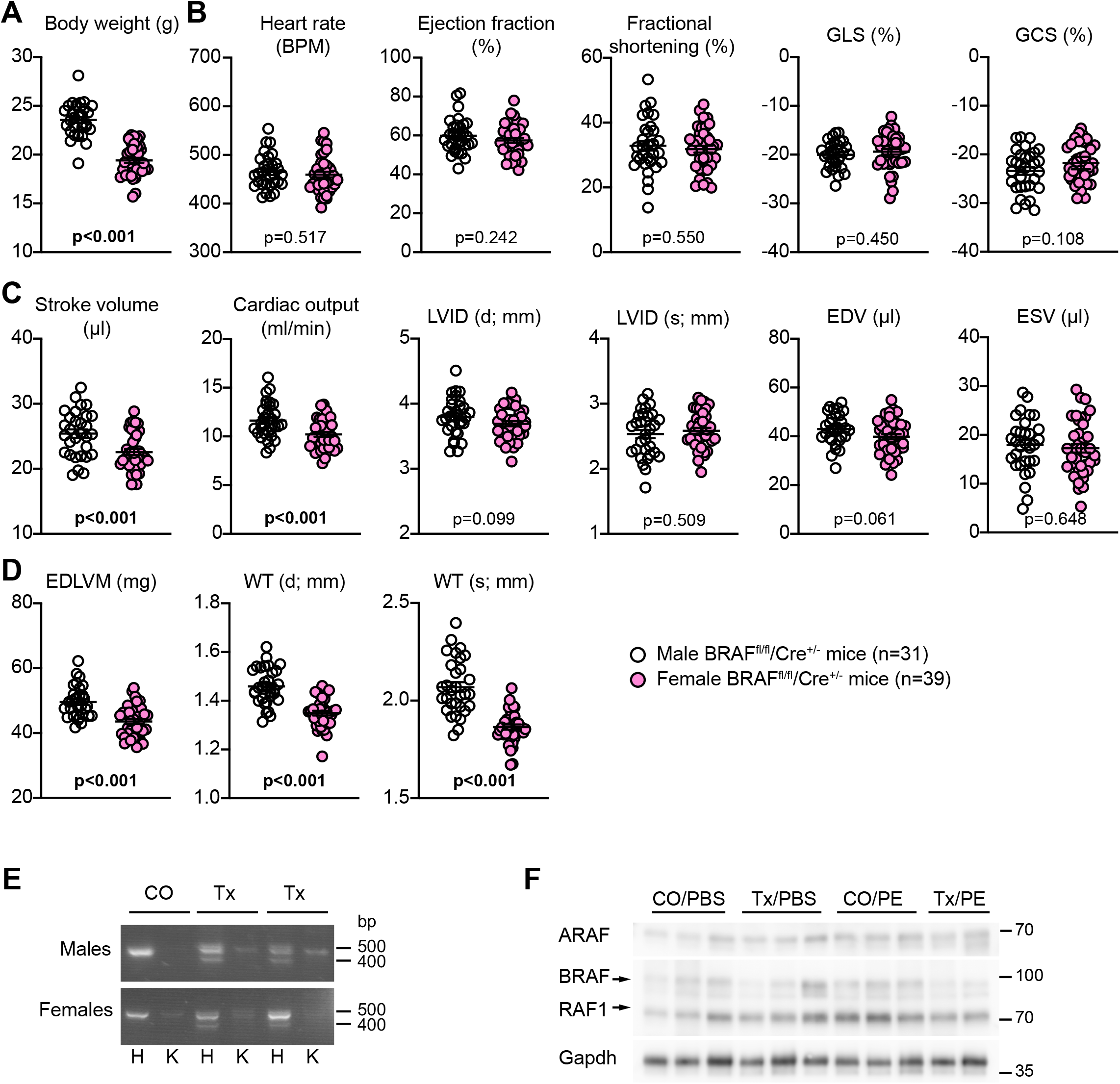
Comparison of cardiac function/dimensions and confirmation of recombination in female BRAF^fl/fl^/Cre^+/-^ mice compared with male littermates. **A-D**, Data were gathered from mice at the time of the first baseline echocardiogram taken from male and female BRAF^fl/fl^/Cre^+/-^ littermates (males: 7-8 weeks; females: 9-10 weeks). **A**, Body weights. **B**, Heart rate, ejection fraction and fractional shortening, global longitudinal strain (GLS) and global circumferential strain (GCS) were measured from B-mode images of long-axis views using VevoStrain speckle-tracking software. **C**,**D**, Stroke volume, cardiac output, end diastolic volume (EDV), end systolic volume (ESV) and end diastolic LV mass (EDLVM) were measured from B-mode images of long-axis views using VevoStrain speckle-tracking software. Diastolic (d) and systolic (s) left ventricle (LV) internal diameter (ID) and wall thickness (WT: anterior plus posterior walls) measurements were measured from M-mode images of short axis views using VevoLab software. **E**, Confirmation of recombination using cDNA prepared from RNA extracted from the hearts of male (upper image) and female (lower image) littermates 11 d post-tamoxifen treatment. PCR amplification used forward primers in exon 9 with reverse primers in exon 13. Deletion of exon 12 in cardiomyocytes resulted in the appearance of a smaller product in heart (H) but not kidney (K) of mice treated with tamoxifen (Tx) in corn-oil but not corn-oil (CO) alone. Representative images are shown. **F**, Immunoblot analysis of RAF isoforms in samples of female mouse hearts treated with CO or Tx 4 days before administration of phenylephrine (PE) in PBS or PBS alone for 7 d. Representative immunoblots are shown.

Female BRAF^fl/fl^/Cre^+/-^ mice were treated with 40 mg/kg tamoxifen or corn-oil vehicle, and then minipumps were implanted for delivery of 40 mg/kg/d phenylephrine using the same schedule as for male mice (**Figure 2A**). Over 7 d, phenylephrine promoted cardiac hypertrophy in the female mice with a decrease in internal diameter and increase in LV wall thickness as assessed using M-mode imaging of the short axis view of the heart (**Figure 6A; Supplementary Table S6**). Peripheral cardiomyocytes were smaller in female hearts than in male hearts, and the increase in cross-sectional area induced by phenylephrine was significant, but less than that detected in male hearts (**Figures 4B and 6B**). Tamoxifen treatment and BRAF knockout resulted in a small decrease in cardiomyocyte size in the female hearts and the minor increase in cardiomyocyte size induced by phenylephrine was less apparent (**Figure 6B**). Phenylephrine increased interstitial fibrosis in some female hearts, particularly in focal areas, but (unlike male hearts) fibrosis was not enhanced with cardiomyocyte BRAF knockout (**Figure 6C-E**). Also in contrast to male hearts, there was no significant effect on perivascular fibrosis. Overall, the response of female mouse hearts to phenylephrine without or with cardiomyocyte BRAF knockout differed from that of the male counterparts.

**Figure 6.**
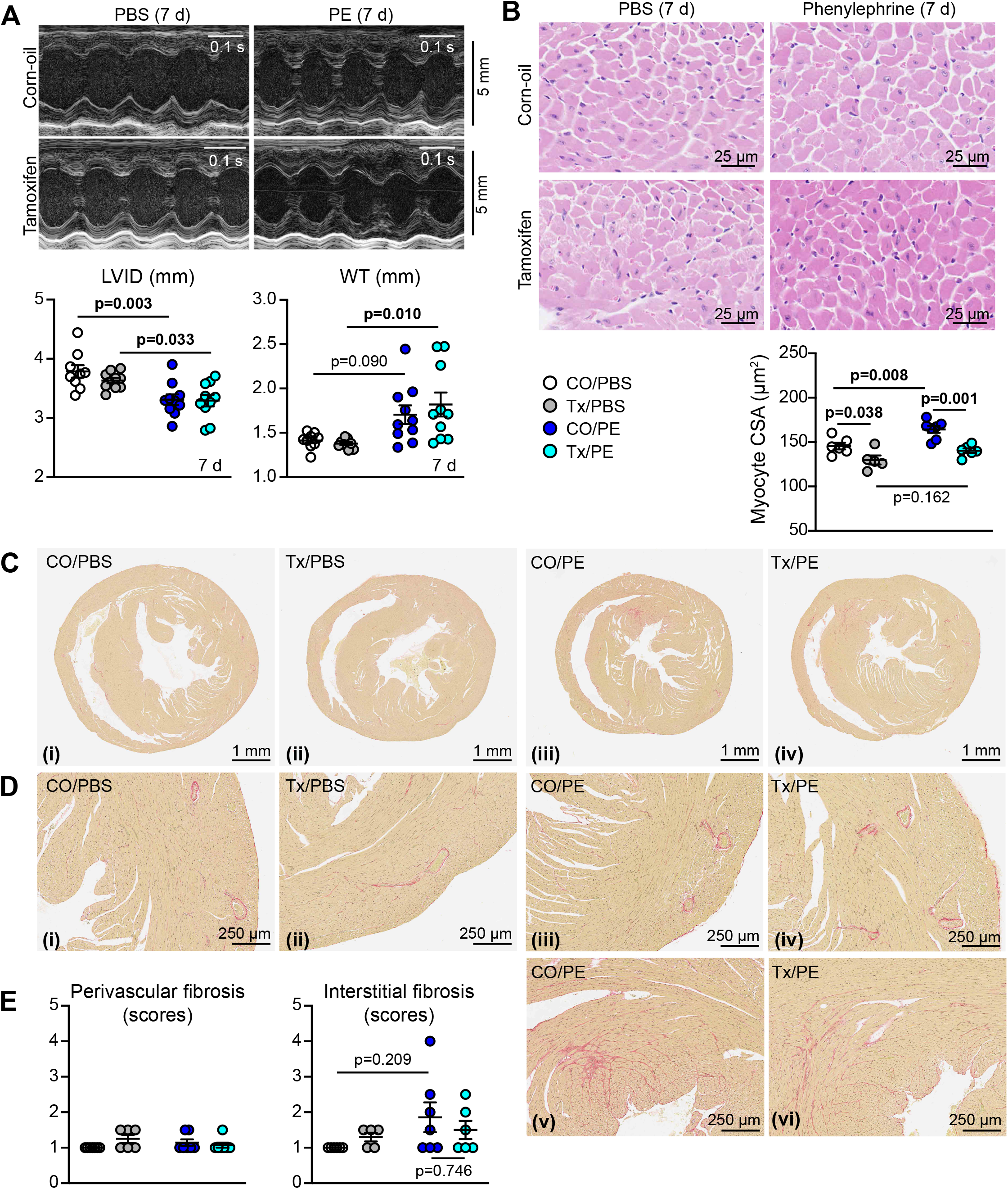
Assessment of the effects of cardiomyocyte BRAF knockout on the response of female mouse hearts to phenylephrine. Female BRAF^fl/fl^/Cre^+/-^ mice were treated with corn-oil (CO) or tamoxifen in corn-oil (Tx) 4 days before minipumps were implanted to deliver PBS or 40 mg/kg/d phenylephrine in PBS (PE) for 7 d. **A**, Representative M-mode echocardiograms taken from short axis views of the heart (upper panels) with analysis of echocardiograms to assess cardiac dimensions (lower panels). LVID, left ventricle internal diameter; WT, left ventricle wall thickness (posterior plus anterior walls). Diastolic measurements are shown. **B**, Haematoxylin and eosin staining of mouse heart sections (upper panels) with assessment of cardiomyocyte cross-sectional area (CSA; lower panel). Images and measurements are from the periphery of the left ventricle. **C**,**D**, Picrosirius red staining of mouse heart sections showing short axis views of the whole heart (**C**) with enlarged sections (**D**) from the same views (the red stain shows accumulation of fibrotic material). Representative (average) images are shown for mice treated with CO/PBS (i), Tx/PBS (ii), CO/PE (iii) and Tx/PE (iv). Additional images are shown for the most severe degree of fibrosis with CO/PE (v) and Tx/PE (vi). **E**, Quantification of fibrosis. This was scored as: 1 = the least amount of fibrosis; 2 = low level fibrosis; 3 = high level fibrosis in at least one area of the myocardium; 4 = high level fibrosis throughout the myocardium (half scores were used). Data are presented as individual values with means ± SEM. Statistical analysis used two-way ANOVA with Holm-Sidak’s post-test. Statistically significant values (p<0.05) are in bold type.

Increasing interstitial fibrosis causes stiffening of the myocardium which leads to diastolic dysfunction. These changes are not necessarily detected using standard M-mode or B-mode echocardiography. Since strain analysis is potentially more sensitive [41], we analysed B-mode images from the phenylephrine study using speckle-tracking software, comparing hearts from male and female mice (**Figure 7A,B**, representative images shown for 7 d only; **Supplementary Tables S5** and **S6**). This approach detected greater baseline variation between animals, so we compared the data obtained at 7 d with baseline data for each mouse individually using paired statistical testing. Heart rate and stroke volumes were unaffected by phenylephrine treatment with or without cardiomyocyte BRAF knockout (**Figure 7C,D**). Consistent with M-mode imaging, phenylephrine promoted a decrease in end diastolic volume and increased end diastolic LV mass in males and females, an effect that was not affected by cardiomyocyte BRAF knockout (**Figure 7E,F**). As with M-mode imaging, there was greater variation in the response of the females and the overall increase in LV mass was relatively less than in males. The lesser degree of hypertrophy induced in female hearts by phenylephrine became apparent in measures of ejection fraction and fractional shortening, in addition to global longitudinal and circumferential strain, all of which were increased by phenylephrine in males, but not females (**Figure 7G-J**). In the males, significant changes were not induced by phenylephrine in hearts with cardiomyocyte BRAF knockout (**Figure 7G,I**), consistent with the adaptive response being compromised with respect to cardiac function, presumably a result of the increase in interstitial fibrosis (**Figure 4C,D**). Surprisingly, in female mice with phenylephrine, cardiomyocyte BRAF knockout resulted in an increase in ejection fraction and fractional shortening, along with increased circumferential strain (**Figure 7H,J**). The reasons are not clear, but the data indicate that BRAF signalling has a different effect in female myocytes than in male myocytes and loss of cardiomyocyte BRAF enhances contractile function.

**Figure 7.**
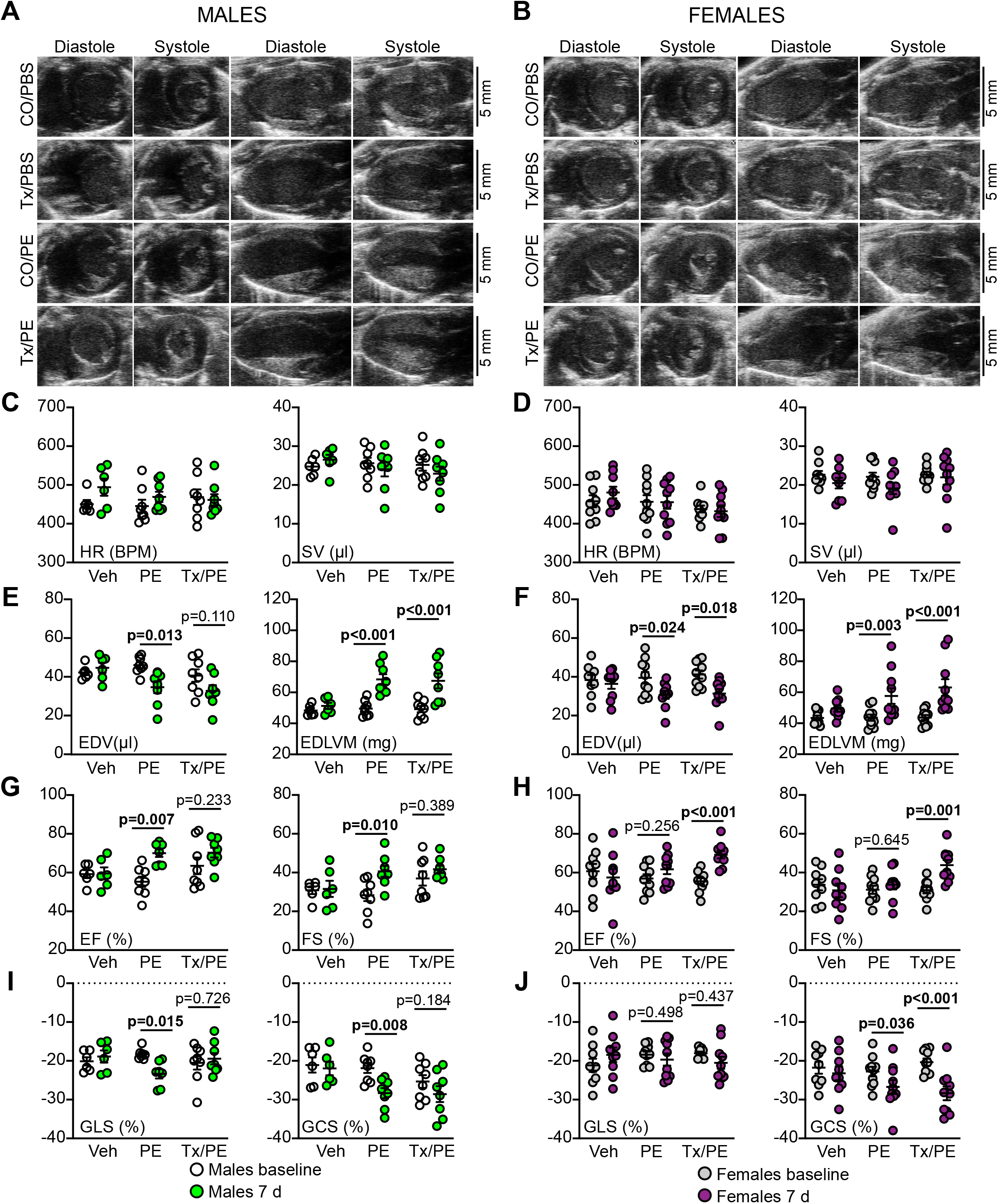
Comparison of effects of cardiomyocyte BRAF knockout on cardiac function in male and female hearts. Male and female BRAF^fl/fl^/Cre^+/-^ mice were treated with corn-oil (CO) or tamoxifen in corn-oil (Tx) 4 days before minipumps were implanted to deliver PBS (PBS) or 40 mg/kg/d phenylephrine in PBS (PE) for 7 d. **A**,**B**, Representative images are shown for short axis (left two panels) and long axis (right two panels) views in diastole or systole in male (**A**) or female mice (**B**). **C-J**, B-mode images were analysed with VevoStrain speckle-tracking software. **C**,**D**, Heart rate (HR) and stroke volume (SV). **E**,**F**, End diastolic volume (EDV) and end diastolic left ventricle mass (EDLVM). **G**,**H**, Ejection fraction (EF) and fractional shortening (FS). **I**,**J**, Global longitudinal strain (GLS) and global circumferential strain (GCS). **C**,**E**,**G**,**I**, Data for male mice; **D**,**F**,**H**,**J**, Data for female mice. All parameters except GCS were measured from long axis views; GCS was taken from short axis views. Data are individual values with means ± SEM. Statistical analysis used paired two-way ANOVA with Holm-Sidak’s post-test. Statistically significant values (p<0.05) are in bold type.

## DISCUSSION

Cardiac hypertrophy is generally considered a predisposing risk factor for heart failure, but cardiomyocyte hypertrophy is a necessary and important adaptation that allows the adult heart to accommodate any increase in workload whether physiological as with exercise or pregnancy, or pathological as with hypertension. Here, we have studied the early adaptive phase, particularly focusing on the response of male mouse hearts to hypertension compared with a more physiological form of hypertrophy induced by phenylephrine. Both cause hypertrophy with reduced LV internal diameter and increased LV wall thickness and mass, but the aetiology is quite different, along with the influence of cardiomyocyte BRAF. In hypertension, even though cardiomyocytes may themselves possess mechanosensors such as stretch-regulated ion channels [30], the primary effect is on the arterioles and the endothelial cells lining the blood vessels in the heart. This was clearly seen in the histology, with a striking and dominant effect of AngII on perivascular fibrosis around the arterioles (**Figure 3A-C**). It is likely that endothelial or smooth muscle cells in the walls of the arterioles produce neurohumoral factors (e.g. endothelin-1 [32]) which then stimulate cardiomyocyte hypertrophy. These act through the various signalling pathways including the ERK1/2 cascade to promote changes in gene and protein expression, including increased expression of growth factors. Our data show that AngII promoted upregulation of *Edn1* and *Fgf2* (as an example of a pro-fibrotic growth factor), with the latter inhibited by cardiomyocyte BRAF knockout suggesting it (but not *Edn1*) originated in the cardiomyocyte (**Figures 2F** and **3D**). FGF2 is likely to stimulate cardiac non-myocytes including resident fibroblasts, and factors such as this are probably responsible for the increase in interstitial fibrosis in this model (**Figure 8A**). In this scenario, loss of BRAF signalling is predicted to reduce cardiomyocyte hypertrophy and interstitial fibrosis as, indeed, it does.

**Figure 8.**
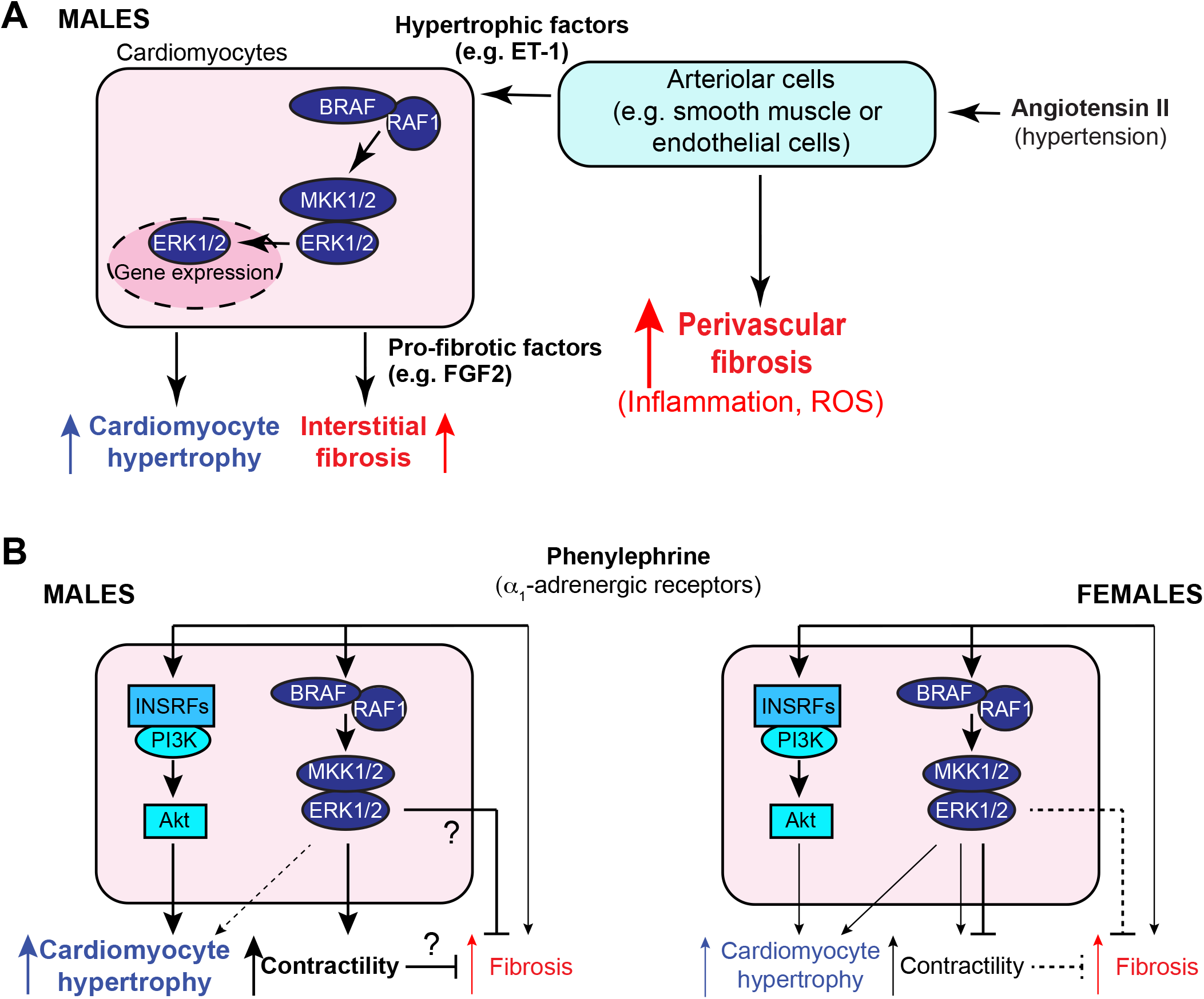
Schematic representations of the conclusions from this study. **A**, Angiotensin II (AngII) causes hypertension and directly stimulates cells in the walls of cardiac arterioles (e.g. endothelial cells or smooth muscle cells). Acutely (over 7 d), this resulted in perivascular fibrosis around arterioles, along with markers of inflammation and increased reactive oxygen species (ROS). These cells also produced hypertrophic factors such as endothelin-1 (ET-1) that stimulate cardiomyocyte hypertrophy via BRAF/RAF1, MKK1/2 and ERK1/2. Cardiomyocyte hypertrophy was associated with production of profibrotic factors such as fibroblast growth factor 2 (FGF2) downstream of BRAF/RAF1→ERK1/2 signalling, and these increased interstitial fibrosis, probably acting on resident fibroblasts. Loss of BRAF resulted in decreased cardiomyocyte hypertrophy and interstitial, but not perivascular, fibrosis. **B**, Phenylephrine acts directly on cardiomyocytes in the heart, but has some additional systemic effects that may lead to limited cardiac fibrosis. In male mice (left), phenylephrine causes cardiomyocyte hypertrophy acting primarily via insulin receptor family members (INSRFs) and Akt (as described in [27]). BRAF/RAF1→ERK1/2 signalling increased contractility and this increase was lost with cardiomyocyte BRAF knockout. In addition, cardiomyocyte BRAF knockout resulted in increased fibrosis, possibly due to loss of a direct inhibitory signal or because of an imbalance between hypertrophy and contractility. In female mice (right), phenylephrine had a modest effect on cardiomyocyte hypertrophy, possibly with some signal from the ERK1/2 cascade, but did not significantly affect contractility. Cardiomyocyte BRAF knockout increased contractility, possibly due to loss of an inhibitory signal, but there was no effect on fibrosis.

As an α_1_-AR agonist phenylephrine acts directly on cardiomyocytes to promote hypertrophy (**Figure 8B**), but the signal to hypertrophy is propagated by transactivation of one or more of the insulin receptor family, signalling via PI3K and PKB/Akt rather than ERK1/2 [27]. Consistent with this, cardiomyocyte BRAF knockout did not significantly affect the increase in cardiomyocyte cross-sectional area induced in male mouse hearts by phenylephrine (**Figures 4B and 8B**). Phenylephrine increased ejection fraction which may be partly attributed to the change in dimensions, but also increased cardiac contractility with increased fractional shortening and longitudinal/circumferential strain, an effect which was lost with cardiomyocyte BRAF knockout (**Figure 7G,I**). Thus, although BRAF signalling may not drive the increase in size in response to phenylephrine, it still influences cardiomyocyte function. This may be mediated by changes in gene expression, but could result from non-genomic effects of ERK1/2 signalling on ion fluxes in the cell. For example, the sodium proton exchanger NHE1 is phosphorylated by p90 ribosomal S6 kinases, downstream of BRAF and the ERK1/2 cascade. This alters the activation profile of NHE1 and is associated with increased contractility [42, 43]. Notably, expression of activated NHE1 in mice enhances the degree of hypertrophy induced by phenylephrine [44] suggesting that it is a contributing factor to the overall response. The other effect of cardiomyocyte BRAF knockout in male mice was to increase cardiac fibrosis, particularly interstitial fibrosis (**Figure 4C-E**). The mechanism is not clear, but could derive from loss of a BRAF signal to inhibit fibrosis, possibly an element of the known cytoprotective aspect of ERK1/2 signalling. Alternatively, it could result from the imbalance resulting from sustained cardiomyocyte hypertrophic growth in a context of compromised contractility. On the other hand, increased fibrosis may be expected to compromise contractility but sustain cardiomyocyte hypertrophy. This interplay between the pro-hypertrophic signalling from PI3K-PKB/Akt and the ERK1/2 cascade clearly requires further study.

Overall, BRAF emerged as a key signalling intermediate in male mice, in the development of both pathological hypertrophy where it has a direct effect on cardiomyocyte size and in physiological hypertrophy to prevent accumulation of fibrotic material during remodelling of the heart. The role of BRAF in cardiomyocytes differs from RAF1 which is potently cytoprotective [19]. This may appear anomalous since our data also show that BRAF and RAF1 exist as preformed heterodimers in cardiomyocytes [16] and they might be expected to have the same role. However, not all RAF1 associates with BRAF, and RAF1 phosphorylates and inhibits pro-apoptotic kinases [16, 45, 46]. There seem to be no reports of the roles of other MKK1/2 activating kinases, ARAF and Tpl2/Cot, in the heart although both are expressed in cardiomyocytes [18]. ARAF may be a partner for RAF1 in cytoprotection, whilst Tpl2/Cot is probably important in the inflammatory response, but further studies could reveal additional specific roles for these kinases.

Despite the increasing appreciation of the importance of heart failure in women, there are still relatively few preclinical studies comparing the responses of male and female hearts. This study highlights the importance of research in this area. Even in the absence of any intervention, there were clear differences in cardiac dimensions that were not simply due to the difference in body weight (**Figure 5A-D**). The smaller LV internal diameter and LV volume resulted in a significant decrease in stroke volume and cardiac output, and the myocardial walls were significantly thinner. Despite this, although measured under anaesthetic, heart rates, ejection fraction and fractional shortening were similar in males and females. Apart from differences at baseline, female hearts exhibited a different response to phenylephrine (**Figure 8B**), and the degree of hypertrophy, though significant, was less pronounced (**Figure 6**). The functional effects were also different, with little or no increase in contractility (**Figure 7H,J**). We currently have no explanation for these differences given the relative paucity of data on female mouse heart function.

Our model (like many) used a system for inducible gene manipulation using a form of Cre that is activated by tamoxifen. In contrast to the male hearts, tamoxifen treatment had a small, but significant inhibitory effect on cardiomyocyte size (**Figure 6B**). Even though the tamoxifen should have cleared the body within 2-3 d of administration [29], the pharmacokinetics in females and in this particular line of mice may differ and there could still be residual effects at the end of the experiment (11 d post-injection). Since tamoxifen is an antagonist for oestrogen and oestrogen promotes PKB/Akt signalling in female hearts [47, 48], it is feasible that the decrease in cardiomyocyte cross-sectional area is due entirely to the effects of tamoxifen. However, it is equally likely that cardiomyocyte BRAF knockout caused the reduction in cardiomyocyte cross-sectional area. Further studies will be necessary to dissect this, probably in parallel with the role of the PI3K→PKB/Akt pathway and insulin receptor family signalling that plays a significant role in male hearts [27]. Perhaps it should also be considered that a different type of cell-specific and inducible genetic system is required for females that does not use tamoxifen and which can also be used in males.

The implications of this study extend beyond just the understanding of the role of cardiomyocyte BRAF in the two models of hypertrophy shown. Not least, our study raises questions about the development of hypertrophy and heart failure in females compared with males, along with the most appropriate systems for studying gene function in females. Clearly, independent and thorough investigation of the responses of the female heart and the roles of individual genes are urgently needed. Nevertheless, BRAF remains a significant therapeutic target for cancer [49] and our data show that it is an important regulator of cardiac hypertrophy at least in male mice. Because existing drugs can activate RAF signalling in cancer cells through the RAF “paradox” [12], other types of inhibitor are in development for more robust inhibition of RAF kinases (e.g. Type 2 inhibitors and BRAF-PROTAC degraders that target BRAF to the proteosome) [49]. Such drugs seem unlikely to have overt on-target cardiotoxic effects in patients without cardiovascular complications (since cardiomyocyte BRAF knockout alone did not affect cardiac function and dimensions), although there may be some risk for patients with underlying cardiac hypertrophy that requires BRAF signalling. On the other hand, BRAF inhibitors may be beneficial in, for example, hypertensive heart disease to reduce interstitial fibrosis. Further studies of cross-talk between different cardiac cell types may provide additional insight and identify new therapeutic approaches to manage fibrosis in the heart.

## Acknowledgements

We thank Andrew Cripps, Mhairi Baxter and Wayne Knight (University of Reading), and Robert Bond, Emma Mustafa and Rene Ocho (St. George’s University of London) for support for the *in vivo* mouse studies.

## Author contributions

H.O.A. was responsible for and conducted the *in vivo* experiments with phenylephrine and assisted with the development of the manuscript. M.A.H. established the cardiomyocyte BRAF knockout mouse line, was responsible for and conducted the *in vivo* experiments with angiotensin II. J.J.C. assisted with the *in vivo* angiotensin II and phenylephrine experiments.T.M. performed the immunoblotting studies for the BRAF knockout mouse line and assisted with the angiotensin II experiments. S.T.E.C. assisted with the *in vivo* experiments with phenylephrine. P.E.G. advised on analysis of mouse echocardiograms. S.J.F. assisted with the angiotensin II studies and with experimental design. P.H.S. contributed to the initiation of the studies, assisted with experimental design and assisted with writing and reviewing the manuscript. A.C. initiated the project, obtained funding, designed the experiments and wrote the manuscript.

## Data Availability Statement

All primary data are available from the corresponding author upon reasonable request. Additional data sharing information is not applicable to this study.

