## Supplementary Tables for "Cardiomyocyte BRAF is a key signalling intermediate in cardiac hypertrophy in mice"

**Supplementary Table S1. Mouse weights.**

**Supplementary Table S2. Primers for genotyping and confirmation of recombination.**

**Supplementary Table S3. qPCR primers.**

**Supplementary Table S4. Echocardiography data for male BRAFKO mice treated without/with angiotensin II.**

**Supplementary Table S5. Echocardiography data for male BRAFKO mice treated without/with phenylephrine.**

**Supplementary Table S6. Echocardiography data for female BRAFKO mice treated without/with phenylephrine.**

**Supplementary Table S1. Mouse weights (g).** Mice were allocated to groups on a random basis and were treated with corn-oil (CO) or tamoxifen in corn-oil (Tx) with acidified PBS (AcPBS) or 0.8 mg/kg/d angiotensin II (AngII), or with PBS or 40 mg/kg/d phenylephrine in PBS (PE). Weights were taken at the start of the study with the first baseline echocardiogram (Start), immediately after minipump surgery, and when mice were culled (End). Weights post-surgery and at the end included the minipumps. Male mice were 7-8 weeks at the start of the experiment; female mice were 9-10 weeks at the start of the experiment.

| **Study** | **Condition** | **Start** | | **Post-minipump** | | **End** | |  |
| --- | --- | --- | --- | --- | --- | --- | --- | --- |
|  |  | **Mean** | **SEM** | **Mean** | **SEM** | **Mean** | **SEM** | **n** |
| AngII (males) | CO/AcPBS | 24.01 | 0.56 | 25.01 | 0.48 | 25.10 | 0.44 | 10 |
| (echocardiography | Tx/AcPBS | 24.70 | 0.48 | 25.93 | 0.44 | 26.13 | 0.40 | 8 |
| + biochemistry) | CO/AngII | 25.17 | 0.38 | 26.60 | 0.33 | 26.12 | 0.64 | 10 |
|  | Tx/AngII | 23.83 | 0.54 | 25.10 | 0.45 | 24.93 | 0.37 | 10 |
| AngII (males) | CO/AcPBS | 23.65 | 0.54 | 26.36 | 0.49 | 26.96 | 0.57 | 8 |
| (histology) | Tx/AcPBS | 22.33 | 0.31 | 25.23 | 0.30 | 26.05 | 0.17 | 4 |
|  | CO/AngII | 24.29 | 0.28 | 26.60 | 0.26 | 26.80 | 0.37 | 7 |
|  | Tx/AngII | 22.88 | 0.70 | 26.30 | 0.56 | 26.36 | 0.58 | 5 |
| PE (males) | CO/PBS | 23.77 | 0.55 | 25.07 | 0.36 | 25.54 | 0.39 | 7 |
|  | Tx/PBS | 23.55 | 0.41 | 25.49 | 0.38 | 25.71 | 0.43 | 8 |
|  | CO/PE | 23.89 | 0.75 | 25.13 | 0.63 | 24.73 | 0.61 | 8 |
|  | Tx/PE | 23.06 | 0.75 | 24.29 | 0.53 | 24.51 | 0.44 | 8 |
| PE (females) | CO/PBS | 19.64 | 0.52 | 21.10 | 0.58 | 21.31 | 0.58 | 9 |
|  | Tx/PBS | 19.19 | 0.39 | 20.96 | 0.28 | 20.99 | 0.21 | 10 |
|  | CO/PE | 19.19 | 0.58 | 20.20 | 0.49 | 20.47 | 0.43 | 10 |
|  | Tx/PE | 19.64 | 0.60 | 20.77 | 0.40 | 20.74 | 0.35 | 10 |

**Supplementary Table S2. Primers for genotyping and confirmation of recombination.**

| **Mouse strain** | **DNA** | **Forward primer** | **Reverse primer** | **Annealing temp.** |
| --- | --- | --- | --- | --- |
| **Genotyping** |  |  |  |  |
| BRAF^fl/fl^ | gDNA | GCATAGCGCATATGCTCACA | CCATGCTCTAACTAGTGCTG | 57°C |
| Cre^-^ | gDNA | TCTATTGCACACAGCAATCCA | CCAACTCTTGTGAGAGGAGCA | 52°C |
| Cre^+^ | gDNA | TCTATTGCACACAGCAATCCA | CCAGCATTGTGAGAACAAGG | 52°C |
| **Recombination** |  |  |  |  |
| BRAF^fl/fl^  (exons 9-13) | cDNA | TTGATTTTGAGCCTGGCCCAGTG | GTGCAGTCTGCCGAGCAATATC | 57°C |
| BRAF^fl/fl^  (exons 10-13) | cDNA | GTCATCTTCTTCCTCATCCTCG | GTGCAGTCTGCCGAGCAATATC | 57°C |

**Supplementary S3. qPCR primers.**

| **Gene Symbol** | **Accession No.** | **Sense Primer (5’→3’)** | **Antisense Primer (5’→3’)** |
| --- | --- | --- | --- |
| Col1a1 | NM_007742 | TCGTGGCTTCTCTGGTCTC | CCGTTGAGTCCGTCTTTGC |
| Col4a1 | NM_009931.2 | TGTGGGCCAGCCAGGCATTG | CAGGGGGTCCGATCGCTCCA |
| Ddr2 | NM_022563.2 | GCACTTGGTGAATTAATTAGAATCCTG | GGACAACTAAATGGTCCCTCCC |
| Edn1 | NM_007913 | GCCTTCGCTCACTCCACTA | GCTGGGATTGGTAGGTGGTA |
| Fn1 | NM_010233 | AAGAGGACGTTGCAGAGCTA | AGACACTGGAGACACTGACTAA |
| Gapdh | NM_008084.2 | TCACCACCATGGAGAAGGC | GCTAAGCAGTTGGTGGTGCA |
| Hif1a | NM_010431 | GATGTAATGTTTCCCTCTTCTAATGA | GCAGGATCAGCACTACTTCG |
| Il6 | NM_031168 | TCCATCCAGTTGCCTTCTTG | GGTCTGTTGGGAGTGGTATC |
| Il1b | NM_008361 | CAACCAACAAGTGATATTCTCCAT | GGGTGTGCCGTCTTTCATTA |
| Lox |  | GACATTCGCTACACAGGACAT | AACACCAGGTACGGCTTTATC |
| Myh6 | NM_010856 | CGAGCTGGATGAGGCGGAG | TCTGCTGGAGAGGTTATTCCTCG |
| Myh7 | NM_080728 | CATGCCAACCGTATGGCTG | GTTCCACGATGGCGATGTTC |
| Nppa | NM_008725 | GATGGATTTCAAGAACCTGCTAGA | CTTCCTCAGTCTGCTCACTCA |
| Nppb | NM_008726 | TCCAGCAGAGACCTCAAAATTC | CAGTGCGTTACAGCCCAAA |
| Postn | NM_015784 | TTCCTCTCCTGCCCTTATATGC | CCTGATCCCGACCCCTGAT |
| NOX1 | NM_172203 \| GCAGGATCAGCACTACTTCG | CATCCCTTCACTCTGACTTCTG | TGCTGTTGTTCAAATGTCCTTATG |
| NOX2 (Cybb) | NM_007807 | GCTATGAGGTGGTGATGTTAGT | GTTTCAGACTGGTGGCATTATC |
| NOX4 | NM_015760 | GGAAGCCCATTTGAGGAGTC | TCCAGTCATCCAGTAGAGTGTT |

**Supplementary Table S4. Echocardiography data for male mice with cardiomyocyte BRAF knockout treated without/with angiotensin II.** Male BRAF^fl/fl^/Cre^+/-^ mice (7-8 weeks) were treated with corn-oil (CO) or tamoxifen in corn-oil (Tx) 4 days before minipumps were implanted for delivery of acidified PBS (AcPBS) or 0.8 mg/kg/d angiotensin II in AcPBS (AngII) for 7 d. Echocardiograms were taken prior to tamoxifen treatment (Baseline) and 7 d after minipump implantation. Echocardiograms were analysed using VevoLab software. LV, left ventricle; ID, internal diameter; AW, anterior wall; PW, posterior wall; WT, wall thickness (AW+PW); EDV, end diastolic volume; ESV, end systolic volume; EDLVM, end diastolic LV mass; ESLVM, end systolic LV mass.

* Data collected from B-mode images of long axis views. Other parameters were measured from M-mode images of short axis views.

| **Baseline** | **CO/AcPBS (n=10)** | | **Tx/AcPBS (n=8)** | | **CO/AngII (n=10)** | | **Tx/AngII (n=10)** | |
| --- | --- | --- | --- | --- | --- | --- | --- | --- |
|  | Mean | SEM | Mean | SEM | Mean | SEM | Mean | SEM |
| Heart Rate (bpm) | 463 | 17 | 476 | 10 | 463 | 10 | 462 | 10 |
| Stroke Volume (µl) | 35.46 | 1.56 | 33.31 | 1.19 | 33.52 | 0.96 | 36.74 | 1.05 |
| Ejection Fraction (%) | 52.30 | 3.72 | 48.47 | 1.88 | 49.01 | 2.51 | 47.92 | 1.31 |
| Fractional Shortening (%) | 27.04 | 2.54 | 24.21 | 1.12 | 24.61 | 1.56 | 23.88 | 0.79 |
| Cardiac Output (ml/min) | 16.26 | 0.64 | 15.85 | 0.68 | 15.46 | 0.51 | 16.95 | 0.57 |
| LVID;s (mm) | 2.929 | 0.163 | 3.020 | 0.093 | 3.016 | 0.128 | 3.165 | 0.043 |
| LVID;d (mm) | 3.984 | 0.110 | 3.978 | 0.076 | 3.985 | 0.097 | 4.157 | 0.031 |
| LVAW;s (mm) | 0.981 | 0.024 | 0.943 | 0.018 | 0.935 | 0.016 | 0.949 | 0.013 |
| LVAW;d (mm) | 0.728 | 0.015 | 0.717 | 0.011 | 0.726 | 0.015 | 0.738 | 0.012 |
| LVPW;s (mm) | 0.967 | 0.036 | 0.909 | 0.032 | 0.939 | 0.016 | 0.941 | 0.028 |
| LVPW;d (mm) | 0.685 | 0.020 | 0.670 | 0.019 | 0.691 | 0.014 | 0.670 | 0.018 |
| EDV (µl)* | 51.36 | 2.91 | 52.97 | 1.85 | 49.45 | 2.35 | 47.74 | 1.56 |
| ESV (µl)* | 26.95 | 2.24 | 28.21 | 1.53 | 27.99 | 2.20 | 25.19 | 1.30 |
| EDLVM (mg)* | 55.87 | 2.19 | 53.42 | 1.77 | 56.67 | 2.28 | 52.32 | 1.96 |
| ESLVM (mg)* | 58.39 | 2.02 | 56.68 | 1.75 | 58.87 | 2.11 | 55.19 | 1.96 |
| **7 d** | **CO/AcPBS (n=10)** | | **Tx/AcPBS (n=8)** | | **CO/AngII (n=10)** | | **Tx/AngII (n=10)** | |
|  | Mean | SEM | Mean | SEM | Mean | SEM | Mean | SEM |
| Heart Rate (bpm) | 488 | 20 | 500 | 9 | 512 | 15 | 489 | 21 |
| Stroke Volume (µl) | 34.43 | 2.01 | 36.09 | 1.85 | 30.63 | 1.78 | 33.95 | 1.90 |
| Ejection Fraction (%) | 49.69 | 3.07 | 51.21 | 2.67 | 57.88 | 3.07 | 57.26 | 4.29 |
| Fractional Shortening (%) | 25.18 | 1.93 | 26.00 | 1.69 | 30.46 | 2.28 | 30.42 | 3.09 |
| Cardiac Output (ml/min) | 16.60 | 0.88 | 18.05 | 1.04 | 15.54 | 0.83 | 16.38 | 0.80 |
| LVID;s (mm) | 3.000 | 0.118 | 2.986 | 0.128 | 2.515 | 0.147 | 2.651 | 0.189 |
| LVID;d (mm) | 3.998 | 0.087 | 4.024 | 0.104 | 3.581 | 0.125 | 3.771 | 0.133 |
| LVAW;s (mm) | 0.976 | 0.026 | 0.980 | 0.034 | 1.160 | 0.043 | 1.122 | 0.081 |
| LVAW;d (mm) | 0.751 | 0.020 | 0.715 | 0.015 | 0.907 | 0.030 | 0.867 | 0.043 |
| LVPW;s (mm) | 0.963 | 0.043 | 0.942 | 0.019 | 1.217 | 0.049 | 1.157 | 0.060 |
| LVPW;d (mm) | 0.699 | 0.031 | 0.662 | 0.018 | 0.899 | 0.032 | 0.815 | 0.049 |
| EDV (µl)* | 47.14 | 2.80 | 51.97 | 3.36 | 39.36 | 2.50 | 37.87 | 3.43 |
| ESV (µl)* | 28.22 | 2.43 | 27.17 | 2.03 | 20.96 | 1.74 | 19.66 | 2.72 |
| EDLVM (mg)* | 52.97 | 1.41 | 55.25 | 1.72 | 69.40 | 2.53 | 64.24 | 4.73 |
| ESLVM (mg)* | 55.55 | 1.77 | 59.09 | 1.71 | 72.32 | 2.60 | 67.32 | 5.07 |

**Supplementary Table S5. Echocardiography data for male mice with cardiomyocyte BRAF knockout treated without/with phenylephrine.** Male BRAF^fl/fl^/Cre^+/-^ mice (7-8 weeks) were treated with corn-oil (CO) or tamoxifen in corn-oil (Tx) 4 days before minipumps were implanted for delivery of PBS or 40 mg/kg/d phenylephrine in PBS (PE) for 7 d. Echocardiograms were taken prior to tamoxifen treatment (Baseline) and 7 d after minipump implantation. Echocardiograms were analysed. LV, left ventricle; ID, internal diameter; AW, anterior wall; PW, posterior wall; WT, wall thickness (AW+PW); EDV, end diastolic volume; ESV, end systolic volume; EDLVM, end diastolic LV mass; ESLVM, end systolic LV mass; GLS, global longitudinal strain; GCS, global longitudinal strain.

* Data collected from B-mode images were analysed using VevoStrain software. Other parameters were measured from M-mode images of short axis views using VevoLab software.

| **Baseline** | **CO/PBS (n=7)** | | **Tx/PBS (n=8)** | | **CO/PE (n=8)** | | **Tx/PE (n=8)** | |
| --- | --- | --- | --- | --- | --- | --- | --- | --- |
|  | Mean | SEM | Mean | SEM | Mean | SEM | Mean | SEM |
| Heart Rate (bpm) | 457 | 10 | 469 | 12 | 459 | 12 | 474 | 15 |
| LVID;s (mm) | 2.697 | 0.125 | 2.421 | 0.113 | 2.723 | 0.096 | 2.315 | 0.127 |
| LVID;d (mm) | 3.908 | 0.126 | 3.729 | 0.102 | 3.920 | 0.091 | 3.653 | 0.088 |
| LVAW;s (mm) | 1.014 | 0.016 | 1.025 | 0.018 | 1.012 | 0.024 | 1.054 | 0.030 |
| LVAW;d (mm) | 0.744 | 0.012 | 0.760 | 0.007 | 0.752 | 0.012 | 0.773 | 0.012 |
| LVPW;s (mm) | 1.003 | 0.018 | 1.093 | 0.037 | 1.011 | 0.030 | 1.058 | 0.037 |
| LVPW;d (mm) | 0.686 | 0.024 | 0.711 | 0.015 | 0.701 | 0.019 | 0.704 | 0.020 |
| EDV (µl)* | 46.42 | 4.29 | 42.65 | 2.43 | 46.10 | 1.52 | 40.69 | 3.20 |
| ESV (µl)* | 19.01 | 1.75 | 16.72 | 1.88 | 20.49 | 1.51 | 15.52 | 2.62 |
| EDLVM (mg)* | 50.25 | 2.32 | 49.31 | 1.12 | 49.51 | 1.80 | 49.24 | 1.99 |
| ESLVM (mg)* | 51.80 | 2.52 | 50.72 | 1.27 | 50.67 | 1.79 | 50.09 | 2.30 |
| Stroke Volume (µl)* | 27.41 | 2.77 | 25.94 | 1.31 | 25.61 | 1.40 | 25.17 | 1.47 |
| Ejection Fraction (%)* | 59.43 | 1.75 | 61.25 | 3.04 | 55.27 | 2.64 | 63.52 | 4.28 |
| Fractional Shortening (%)* | 31.23 | 1.70 | 34.86 | 2.59 | 28.28 | 2.97 | 36.95 | 3.67 |
| Cardiac Output (ml/min)* | 12.30 | 1.26 | 12.13 | 0.70 | 11.46 | 0.83 | 11.67 | 0.57 |
| GLS (%)* | -20.62 | 1.02 | -21.27 | 1.02 | -19.03 | 0.88 | -20.52 | 1.72 |
| GCS (%)* | -21.43 | 1.69 | -24.66 | 1.48 | -21.89 | 1.24 | -25.33 | 1.66 |
| **7 d** | **CO/PBS (n=7)** | | **Tx/PBS (n=8)** | | **CO/PE (n=8)** | | **Tx/PE (n=8)** | |
|  | Mean | SEM | Mean | SEM | Mean | SEM | Mean | SEM |
| Heart Rate (bpm) | 485 | 18 | 513 | 9 | 470 | 15 | 465 | 17 |
| LVID;s (mm) | 2.622 | 0.130 | 2.636 | 0.118 | 2.137 | 0.151 | 1.920 | 0.126 |
| LVID;d (mm) | 3.841 | 0.119 | 3.917 | 0.105 | 3.505 | 0.121 | 3.307 | 0.077 |
| LVAW;s (mm) | 1.041 | 0.028 | 1.101 | 0.031 | 1.276 | 0.064 | 1.297 | 0.076 |
| LVAW;d (mm) | 0.771 | 0.012 | 0.801 | 0.017 | 0.865 | 0.038 | 0.917 | 0.054 |
| LVPW;s (mm) | 1.029 | 0.028 | 1.072 | 0.032 | 1.301 | 0.095 | 1.405 | 0.099 |
| LVPW;d (mm) | 0.697 | 0.027 | 0.708 | 0.024 | 0.974 | 0.089 | 1.000 | 0.094 |
| EDV (µl)* | 47.61 | 3.65 | 48.06 | 3.10 | 34.69 | 3.07 | 32.84 | 2.94 |
| ESV (µl)* | 18.74 | 1.97 | 19.72 | 2.10 | 10.64 | 1.45 | 9.91 | 1.47 |
| EDLVM (mg)* | 53.23 | 2.60 | 51.93 | 1.41 | 68.34 | 3.36 | 67.43 | 4.89 |
| ESLVM (mg)* | 55.29 | 2.98 | 53.82 | 1.50 | 70.73 | 3.65 | 69.15 | 5.33 |
| Stroke Volume (µl)* | 28.87 | 2.58 | 28.33 | 1.50 | 24.04 | 1.83 | 22.93 | 1.83 |
| Ejection Fraction (%)* | 60.44 | 2.76 | 59.13 | 2.42 | 70.04 | 1.95 | 70.30 | 2.48 |
| Fractional Shortening (%)* | 32.02 | 3.56 | 32.85 | 2.40 | 40.67 | 2.84 | 41.67 | 1.90 |
| Cardiac Output (ml/min)* | 13.67 | 0.87 | 13.83 | 0.92 | 11.29 | 0.92 | 10.65 | 1.00 |
| GLS (%)* | -19.70 | 1.55 | -20.90 | 1.18 | -23.33 | 1.05 | -19.44 | 1.44 |
| GCS (%)* | -22.79 | 1.69 | -23.22 | 0.75 | -28.33 | 1.33 | -28.57 | 2.05 |

**Supplementary Table S6. Echocardiography data for female mice with cardiomyocyte BRAF knockout treated without/with phenylephrine.** Female BRAF^fl/fl^/Cre^+/-^ mice (9-10 weeks) were treated with corn-oil (CO) or tamoxifen in corn-oil (Tx) 4 days before minipumps were implanted for delivery of PBS or 40 mg/kg/d phenylephrine in PBS (PE) for 7 d. Echocardiograms were taken prior to tamoxifen treatment (Baseline) and 7 d after minipump implantation. Echocardiograms were analysed. LV, left ventricle; ID, internal diameter; AW, anterior wall; PW, posterior wall; WT, wall thickness (AW+PW); EDV, end diastolic volume; ESV, end systolic volume; EDLVM, end diastolic LV mass; ESLVM, end systolic LV mass; GLS, global longitudinal strain; GCS, global longitudinal strain.

* Data collected from B-mode images were analysed using VevoStrain software. Other parameters were measured from M-mode images of short axis views using VevoLab software.

| **Baseline** | **CO/PBS (n=9)** | | **Tx/PBS (n=10)** | | **CO/PE (n=10)** | | **Tx/PE (n=10)** | |
| --- | --- | --- | --- | --- | --- | --- | --- | --- |
|  | Mean | SEM | Mean | SEM | Mean | SEM | Mean | SEM |
| Heart Rate (bpm) | 457 | 17 | 464 | 7 | 465 | 14 | 452 | 10 |
| LVID;s (mm) | 2.576 | 0.104 | 2.556 | 0.061 | 2.569 | 0.113 | 2.636 | 0.089 |
| LVID;d (mm) | 3.726 | 0.067 | 3.632 | 0.055 | 3.671 | 0.113 | 3.743 | 0.069 |
| LVAW;s (mm) | 0.941 | 0.008 | 0.948 | 0.011 | 0.959 | 0.021 | 0.917 | 0.014 |
| LVAW;d (mm) | 0.713 | 0.008 | 0.716 | 0.008 | 0.696 | 0.015 | 0.700 | 0.009 |
| LVPW;s (mm) | 0.938 | 0.026 | 0.917 | 0.012 | 0.907 | 0.018 | 0.933 | 0.018 |
| LVPW;d (mm) | 0.639 | 0.011 | 0.657 | 0.018 | 0.650 | 0.021 | 0.650 | 0.011 |
| EDV (µl)* | 38.34 | 2.64 | 40.18 | 1.15 | 39.53 | 2.96 | 41.20 | 2.10 |
| ESV (µl)* | 15.70 | 2.42 | 17.31 | 1.24 | 17.42 | 2.07 | 18.52 | 1.57 |
| EDLVM (mg)* | 43.46 | 1.48 | 43.45 | 1.21 | 43.65 | 1.99 | 43.69 | 1.57 |
| ESLVM (mg)* | 44.50 | 1.53 | 44.70 | 1.61 | 45.34 | 2.14 | 44.76 | 1.66 |
| Stroke Volume (µl)* | 22.64 | 0.99 | 22.87 | 1.01 | 22.11 | 1.08 | 22.68 | 0.72 |
| Ejection Fraction (%)* | 60.72 | 3.79 | 57.11 | 2.54 | 56.94 | 2.25 | 55.50 | 1.68 |
| Fractional Shortening (%)* | 33.71 | 2.86 | 31.65 | 2.09 | 31.15 | 2.09 | 30.91 | 1.62 |
| Cardiac Output (ml/min)* | 10.34 | 0.50 | 10.49 | 0.56 | 10.11 | 0.62 | 9.93 | 0.34 |
| GLS (%)* | -21.15 | 1.69 | -19.50 | 1.33 | -18.41 | 0.78 | -18.69 | 0.76 |
| GCS (%)* | -21.76 | 1.54 | -22.21 | 1.36 | -22.76 | 1.23 | -20.35 | 0.96 |
| **7 d** | **CO/PBS (n=9)** | | **Tx/PBS (n=10)** | | **CO/PE (n=10)** | | **Tx/PE (n=10)** | |
|  | Mean | SEM | Mean | SEM | Mean | SEM | Mean | SEM |
| Heart Rate (bpm) | 487 | 16 | 478 | 13 | 471 | 13 | 447 | 11 |
| LVID;s (mm) | 2.570 | 0.142 | 2.508 | 0.093 | 1.995 | 0.118 | 1.946 | 0.109 |
| LVID;d (mm) | 3.786 | 0.106 | 3.632 | 0.045 | 3.310 | 0.090 | 3.293 | 0.098 |
| LVAW;s (mm) | 1.004 | 0.025 | 1.001 | 0.017 | 1.183 | 0.058 | 1.261 | 0.086 |
| LVAW;d (mm) | 0.739 | 0.021 | 0.726 | 0.010 | 0.821 | 0.031 | 0.900 | 0.052 |
| LVPW;s (mm) | 1.004 | 0.045 | 0.958 | 0.021 | 1.249 | 0.076 | 1.282 | 0.096 |
| LVPW;d (mm) | 0.674 | 0.014 | 0.677 | 0.022 | 0.883 | 0.076 | 0.919 | 0.090 |
| EDV (µl)* | 36.36 | 2.52 | 38.51 | 1.78 | 30.70 | 2.06 | 31.39 | 2.33 |
| ESV (µl)* | 15.89 | 2.34 | 15.81 | 1.40 | 11.57 | 1.00 | 9.42 | 0.73 |
| EDLVM (mg)* | 49.56 | 2.09 | 51.12 | 1.66 | 57.60 | 5.03 | 63.09 | 5.35 |
| ESLVM (mg)* | 51.19 | 2.05 | 52.90 | 1.90 | 59.42 | 5.29 | 64.74 | 5.32 |
| Stroke Volume (µl)* | 20.47 | 1.37 | 22.71 | 1.02 | 19.13 | 1.49 | 21.97 | 1.85 |
| Ejection Fraction (%)* | 57.50 | 4.36 | 59.04 | 2.27 | 61.68 | 2.44 | 69.25 | 1.90 |
| Fractional Shortening (%)* | 30.83 | 3.50 | 32.53 | 1.88 | 33.77 | 2.45 | 43.86 | 2.60 |
| Cardiac Output (ml/min)* | 9.91 | 0.85 | 11.01 | 0.67 | 8.64 | 0.69 | 9.65 | 1.03 |
| GLS (%)* | -18.45 | 1.86 | -19.59 | 0.72 | -19.67 | 1.55 | -20.51 | 1.55 |
| GCS (%)* | -23.21 | 1.78 | -21.77 | 1.53 | -26.72 | 1.78 | -28.36 | 1.77 |
